# Spatial mapping of cellular and molecular plasticity in the maternal and postpartum mouse brain

**DOI:** 10.64898/2026.01.03.697464

**Authors:** Keon Arbabi, Shahrzad Ghazisaeidi, Minjin Kim, Bharti Kukreja, Min Yi Feng, Spencer D. Pogue, Hugo Hudson, Maria Eleni Fafouti, Elizabeth E. Crouch, Liisa A.M. Galea, Shreejoy J. Tripathy, Brian T. Kalish

## Abstract

Pregnancy is a critical window for neuroplasticity and maternal mental health, yet our understanding of the molecular and cellular changes underlying this adaptation remains incomplete. Here we profile the female mouse brain in nulliparous, late-pregnant and postpartum states with three single-cell spatial technologies (Slide-tags, MERFISH and Xenium). Together these assays resolve 1.5 million cells in one coronal plane spanning the cortex, striatum, lateral septum and preoptic area. Pregnancy is associated with changes in gene expression in most cell types, beyond the circuits previously established to govern maternal behavior. These changes resolve into three programs that are reproducible across platforms: synaptic pathways increase in neurons as growth and plasticity pathways decrease; immune and angiogenic pathways increase in glia and vascular cells; and cholesterol synthesis decreases as uptake increases across both neurons and glia. Finally, mapping human depression genetics onto these data, we find risk genes concentrated almost entirely in neurons, and this concentration changes with reproductive state in the preoptic area, basal forebrain and ventral striatum. These results place the neurons carrying depression risk among the circuits remodeled during pregnancy, providing a possible cellular substrate for peripartum vulnerability.

## Introduction

The maternal brain remodels during pregnancy to support the onset of parental care, driven by sustained shifts in ovarian and placental hormones^1–3^. In humans, longitudinal imaging has tracked this remodeling at the macroscopic scale, most consistently as a loss of cortical gray-matter volume during gestation that only partly reverses after birth^4–7^. The same window carries a sharp increase in psychiatric risk. Admissions peak in the weeks after delivery^8^, and peripartum depression affects one in six to one in four mothers^9,10^. Thus, maternal adaptation and psychiatric vulnerability coincide, suggesting their cellular substrates may overlap.

Although the macroscopic and behavioral changes of the peripartum period are well described^3,7^, their cellular basis is only partially resolved. The clearest picture is of the medial preoptic area (MPOA), the principal hub for parental behavior^11,12^, where hormone-sensitive neurons remodel in anticipation of motherhood^13^ and neurons that govern parenting and satiety become more active postpartum^14^. Cellular changes have also been resolved in the hippocampus^15,16^ and among the pregnancy-recruited stem cell pools of the ventricular-subventricular zone^17^. Broader surveys have covered more of the brain but only in bulk, one region at a time, in the septum^18,19^, preoptic area^20^, nucleus accumbens^21^ and prefrontal cortex^22^. A recent study spans 11 regions in bulk but resolves cell types in the dorsal hippocampal formation alone^23^. What is missing is a view that spans regions and cell types at once, placing these focal changes alongside the broader neuronal, glial and vascular landscape.

Here we profile the female mouse brain across three reproductive states (nulliparous, late pregnancy and late postpartum) with three complementary single-cell spatial technologies. We leverage Slide-tags, providing whole-transcriptome coverage^24^ alongside MERFISH^25^ and Xenium^26^, imaging 500- and 5,000-gene panels directly in tissue, respectively. Together these assays resolve ∼1.5 million cells in a single coronal plane spanning the isocortex, striatum, lateral septum and MPOA. We annotated every cell against the Allen Institute mouse whole-brain atlas^27^ and combined the platforms by cross-platform meta-analysis, retaining only changes reproducible on at least two technologies. Pregnancy changed gene expression in most cell types without a coordinated change in cellular composition or spatial organization, resolving into neuronal, glial-vascular and lipid-metabolic programs. These programs differed in persistence: some reset after birth, whereas others carried into the postpartum period. Integrating spatial transcriptomics with human GWAS, we found that genetic risk for psychiatric disorders is concentrated in neurons and shifts with reproductive state, offering cell type-resolved hypotheses for peripartum vulnerability.

## Results

### Spatially resolved single-cell transcriptomic profiling of the maternal brain

We studied the maternal mouse brain across three reproductive states: nulliparous (virgin), late pregnancy (gestational day 18), and late postpartum (day 20). These stages span the peripartum window, capturing the hormonal peak of late gestation and the late-lactation state before weaning^1^. For each animal, we sampled a single coronal plane (approximately Bregma +0.2 mm) encompassing the isocortex, striatum, lateral septum, and MPOA (Supplementary Fig. 1). These regions house circuits that govern maternal behavior and undergo pronounced synaptic and structural remodeling across the peripartum period^2,3,13^.

We profiled our coronal sections with three single cell-resolution spatial transcriptomics platforms (Fig. 1A). Slide-tags adds spatial position to whole-transcriptome single-nucleus RNA sequencing (snRNA-seq) by assigning spatial barcodes to nuclei in intact tissue before dissociation^24^. MERFISH and Xenium complement this with single-molecule imaging, detecting individual transcripts of a predefined gene panel directly in the tissue at high spatial resolution ^25,26^. For MERFISH, we designed a custom 500-gene panel focused on cell type markers and pregnancy-relevant pathways from the literature. For Xenium, we employed the Prime 5K Mouse Pan Tissue & Pathways Panel, reflecting a broader 5,000-gene panel that samples the transcriptome beyond a pregnancy-specific focus (Supplementary Data 1) .

**Fig. 1.**
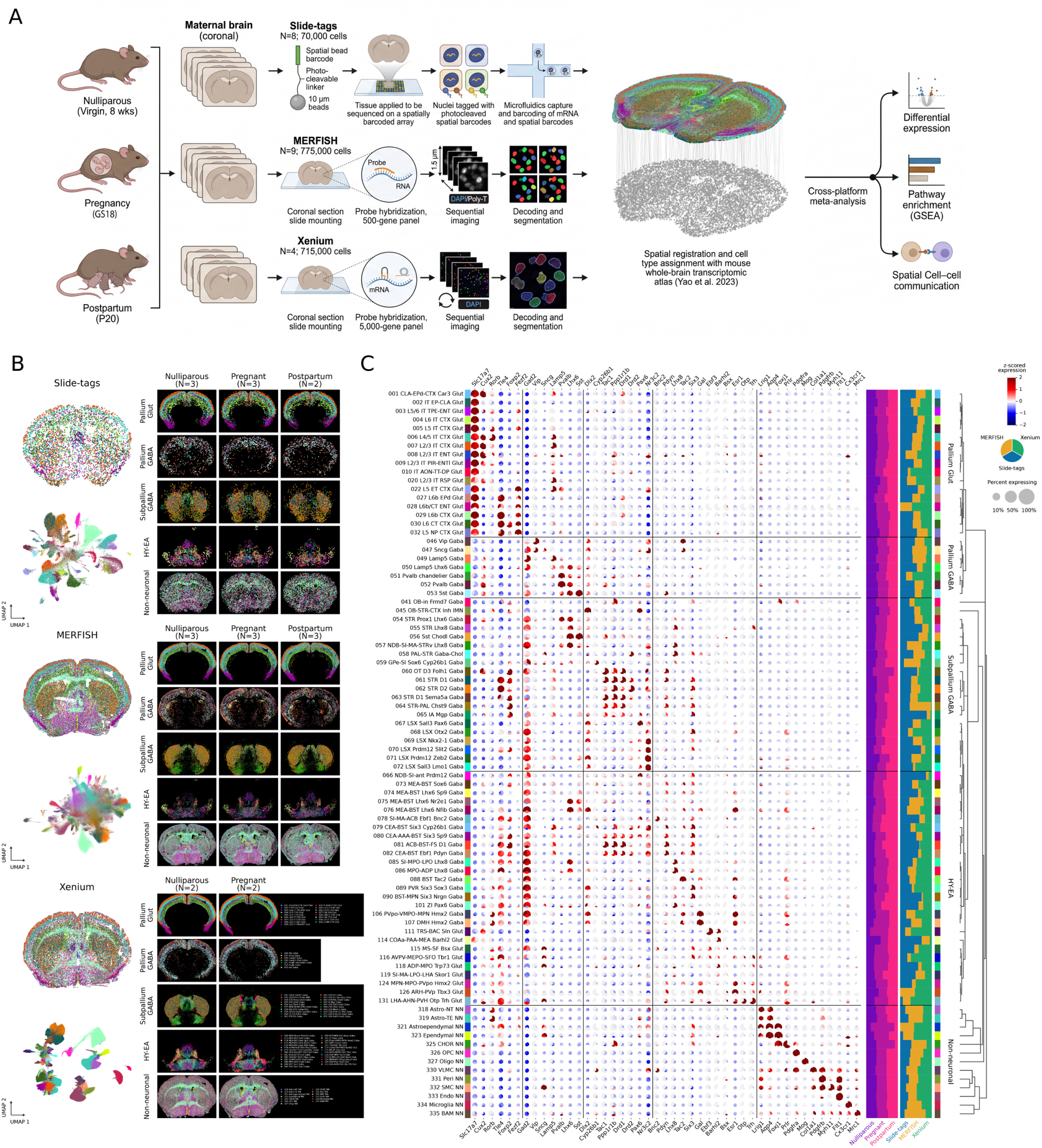
Spatially resolved single-cell transcriptomic profiling of the maternal mouse brain. **A)** Experimental design. Brains from three maternal states – virgin control (8 weeks), pregnant (gestational day 18), and postpartum (day 20) – were profiled with three spatial transcriptomic technologies: Slide-tags (whole transcriptome; *n* = 8, ∼70,000 cells), MERFISH (500-gene panel; *n* = 9, ∼775,000 cells), and Xenium (5,000-gene panel; *n* = 4, ∼715,000 cells). All datasets were spatially registered to the Allen mouse whole-brain atlas^27^ for unified cell type annotation, then integrated by cross-platform meta-analysis for differential expression, pathway enrichment (GSEA), and spatial cell–cell communication. **B)** Annotated cell types for each technology (Slide-tags, top; MERFISH, middle; Xenium, bottom). Left: a representative section with cells colored by annotated cell type (top) and a UMAP of all cells colored by cell type (bottom). Right: spatial distribution of five major cell neighborhoods – pallium glutamatergic, pallium GABAergic, subpallium GABAergic, hypothalamus–extended amygdala (HY-EA), and non-neuronal – across the experimental conditions, with the number of samples per condition indicated (postpartum not profiled by Xenium). **C)** Combined marker-gene dot plot for the 84 cell types (subclasses) robustly detected across platforms (≥100 cells in at least two of the three technologies), grouped by neighborhood and ordered by Allen taxonomy within each. Each dot comprises three wedges, one per technology (Slide-tags, MERFISH, Xenium); wedge radius denotes the fraction of cells expressing the gene and fill color the z-scored mean expression, with gray wedges marking genes absent from a platform’s panel. Marker genes (columns) are grouped by the neighborhood they label. Right: a Xenium-derived dendrogram of transcriptomic similarity, a subclass color key, and stacked bars giving, per cell type, the proportional representation of each condition and of each technology.

Across three separate cohorts, we profiled ∼1.5 million cells: ∼70,000 nuclei (Slide-tags, 8 mice), ∼775,000 cells (MERFISH, 9 mice), and ∼715,000 cells (Xenium, 4 mice) (Supplementary Data 1). We used Xenium to sample the nulliparous and pregnant states, and Slide-tags and MERFISH to additionally sample the postpartum state.

To annotate cell types consistently across platforms, we transferred labels from the Allen mouse whole-brain atlas, which comprises two reference datasets under a single taxonomy: a MERFISH spatial reference that provides each cell type’s anatomical distribution, and a single-cell RNA-sequencing reference (scRNA-seq; 2.3 million cells) that provides its expression signature^27^. Using CAST, a graph neural network for spatial transcriptomics^28^, we registered each query sample to the common coordinate framework (CCFv3) of the spatial reference, enabling a two-step, spatially informed label transfer (see Methods). First, we took the cell types within a physical radius of each query cell as candidates; second, we selected among these by matching the query cell’s expression to the scRNA-seq reference. This approach improved annotation robustness by keeping identity assignments consistent with anatomical location, resolving subclasses that expression alone may confuse (Supplementary Fig. 3). Registration also placed every section on the reference anteroposterior axis, where conditions differed by at most 0.12 mm, less than the median 0.18 mm spacing between adjacent reference sections, so the sampled planes are reasonably well matched (Supplementary Fig. 4).

We resolved 84 cell subclasses, spanning 16 broad classes, each detected in at least two platforms and with expression of established marker genes (Fig. 1C). Neuronal subclasses included layer-specific cortical excitatory neurons (*Slc17a7*), GABAergic interneurons (*Pvalb*, *Sst*, *Vip*, *Lamp5*), striatal medium spiny neurons (*Drd1*, *Drd2*), and preoptic and hypothalamic populations (*Lhx8*, *Gal*, *Esr1*). Non-neuronal subclasses included astrocytes (*Aqp4*), oligodendrocytes and their precursors (*Mog*, *Pdgfra*), microglia (*Cx3cr1*), ependymal cells (*Foxj1*), and vascular cells, including endothelial cells (*Flt1*), pericytes (*Pdgfrb*), and smooth muscle (*Myh11*). Each subclass formed a coherent cluster in each platform’s UMAP and occupied its expected anatomical position (Fig. 1B), with its canonical markers expressed consistently across all three platforms (Fig. 1C; Supplementary Data 1).

We first asked whether pregnancy shifts the composition or spatial arrangement of brain cell types within our sections. The relative proportion of each cell type did not change reproducibly: no subclass reached an FDR <0.10 on any platform, and the few nominal shifts we detected did not agree in direction across platforms. The same held for local organization. Using data from the imaging datasets, for each pair of subclasses, we scored how enriched or depleted one was among the other’s neighbors within ∼200 µm, relative to shuffled cell type labels (see Methods). While some pairs shifted in the context of some datasets, they did not add up to a coherent change: no cell type consistently changed its neighbors, and no group of related cell types shifted together (Supplementary Fig. 5; Supplementary Data 2). We note that modest sample size limits sensitivity to subtle changes in spatial arrangement.

### Pregnancy remodels the maternal brain through distinct neuronal, glial-vascular, and lipid programs

Each spatial platform carries distinct technical biases, from cell segmentation to gene-panel design and detection sensitivity^29^. To distinguish genuine biology from technical artifacts, we first corrected the segmentation and transcript spillover of spatial assays^30,31^ with ResolVi, a generative model that separates each cell’s intrinsic expression from neighbor contamination and background^31^. We then combined per-cell-type results across platforms with SumRank, a published rank-based method for identifying effects that replicate across datasets^32^, keeping only changes reproducible on at least two platforms (D ≥ 2) at an empirical *p* ≤ 0.05 calibrated against permuted condition labels. We applied this framework to differential expression and pathway enrichment, and an analogous concordance criterion to cell–cell communication (see Methods). This threshold is exploratory, and no individual change survives correction across all 89,457 gene and cell type tests (smallest false discovery rate 0.19; Supplementary Data 3). We therefore organize the results by pathway enrichment and report only changes that reproduce on more than one platform.

Pregnancy changed gene expression across most cell types, well beyond the established circuits that are known to govern maternal behavior. We restricted the meta-analysis to the 3,881 genes measured on at least two platforms (see Methods). Of the 84 cell types, 61 carried candidate changes in pregnancy, spanning neuronal and non-neuronal classes (pregnant versus nulliparous; Fig. 2A). Each changed by a median of about 160 genes. The largest shifts occurred in cortical excitatory neurons, striatal medium spiny neurons, endothelial cells and astrocytes (more than 340 genes each; Fig. 2A). The postpartum contrasts drew on Slide-tags and MERFISH alone, because Xenium did not sample the postpartum state, restricting them to roughly 350 shared genes against 3,881 for pregnancy. Their smaller counts therefore reflect the narrower gene space rather than a smaller biological effect, and are not directly comparable with the pregnancy contrast.

**Fig. 2.**
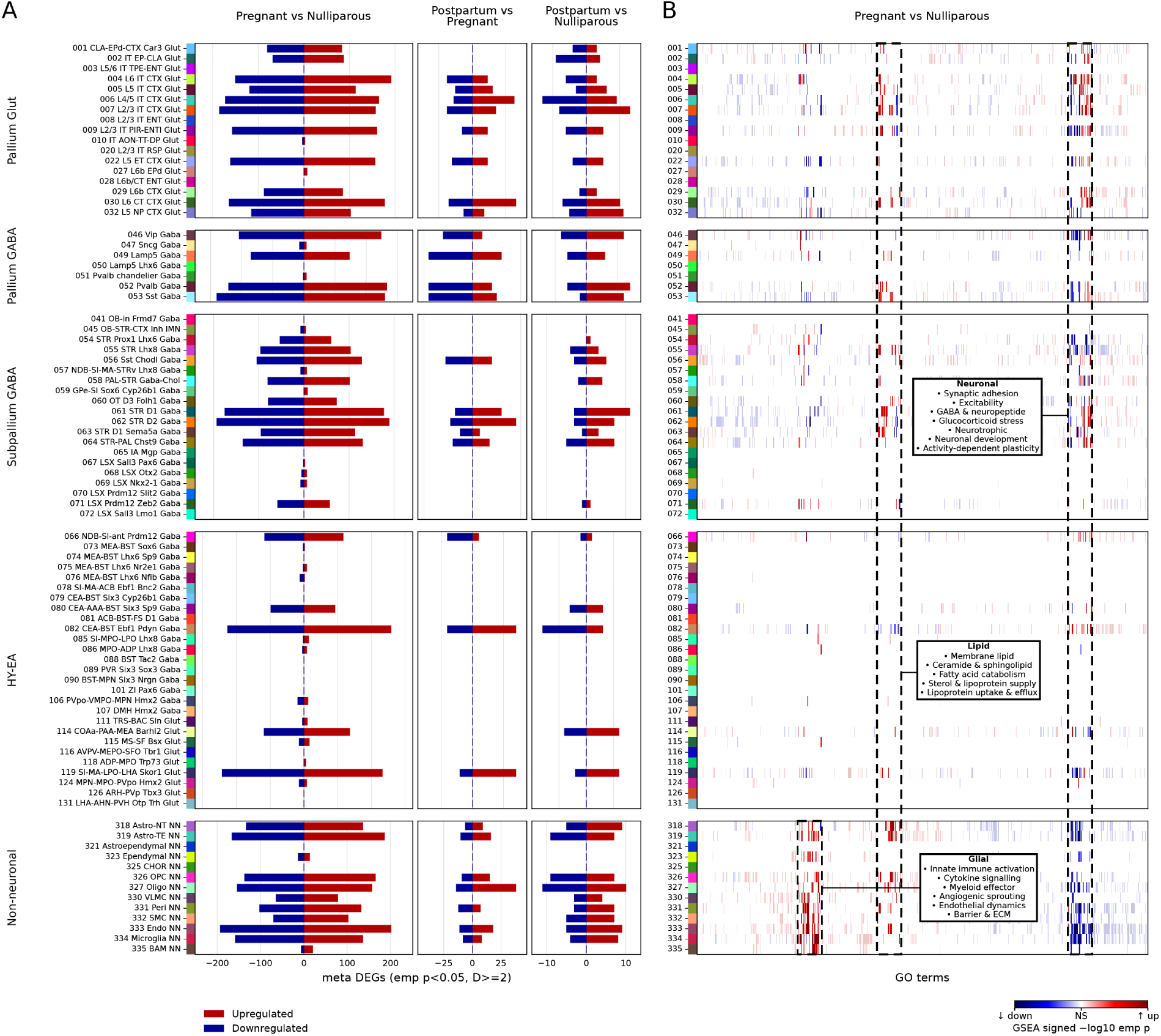
Cell type-resolved meta-analysis of gene expression and pathway enrichment in late pregnancy. Differential expression and gene set enrichment (GSEA) were meta-analyzed across the three spatial technologies, retaining effects reproducible in ≥ 2 of 3 platforms (empirical *p* < 0.05, D ≥ 2). Both panels show the 84 cross-platform subclasses, grouped by neighborhood. **A)** Number of meta-differentially expressed genes per cell type, by direction (upregulated, red; downregulated, blue), for three contrasts: pregnant vs nulliparous, postpartum vs pregnant, and postpartum vs nulliparous. Counts are of genes nominated at the exploratory threshold (empirical *p* ≤ 0.05 with at least two contributing platforms; Methods), not false-discovery-rate-corrected calls. **B)** Pathway enrichment for the pregnant vs nulliparous contrast. Columns are all GO biological-process terms significant in ≥ 1 cell type, ordered by hierarchical clustering of their enrichment profiles. Fill color shows the signed enrichment significance (signed −log10 empirical p; red = up, blue = down in pregnancy; non-significant or untested, white). Curated terms grouped into three super-themes (Neuronal, Glial, and Lipid) are highlighted with their constituent programs.

We grouped the enriched pathways into three programs, each recurring across many cell types (Fig. 2B; Supplementary Data 4). In neurons, pathways for synaptic adhesion, excitability, GABAergic signaling and neuropeptides increased, along with glucocorticoid and neurotrophic signaling, whereas pathways for neuronal development and activity-dependent plasticity decreased. In non-neuronal cells, immune and vascular programs increased, spanning inflammatory and cytokine pathways in glia and angiogenic, endothelial and barrier pathways in the vasculature. A third, lipid-metabolic program spanned neurons and glia: pathways for membrane lipids, sphingolipids and fatty acids increased, and cholesterol provisioning shifted from synthesis and supply toward lipoprotein uptake.

### A transient synaptic and hormonal response resolves into a consolidated neuronal retuning

In pregnancy, synaptic, hormonal and developmental pathways changed in 33 of the 42 neuronal subclasses in which enrichment could be scored on at least two platforms, spanning the cortex, striatum and extended amygdala (Fig. 3B,C).

**Fig. 3.**
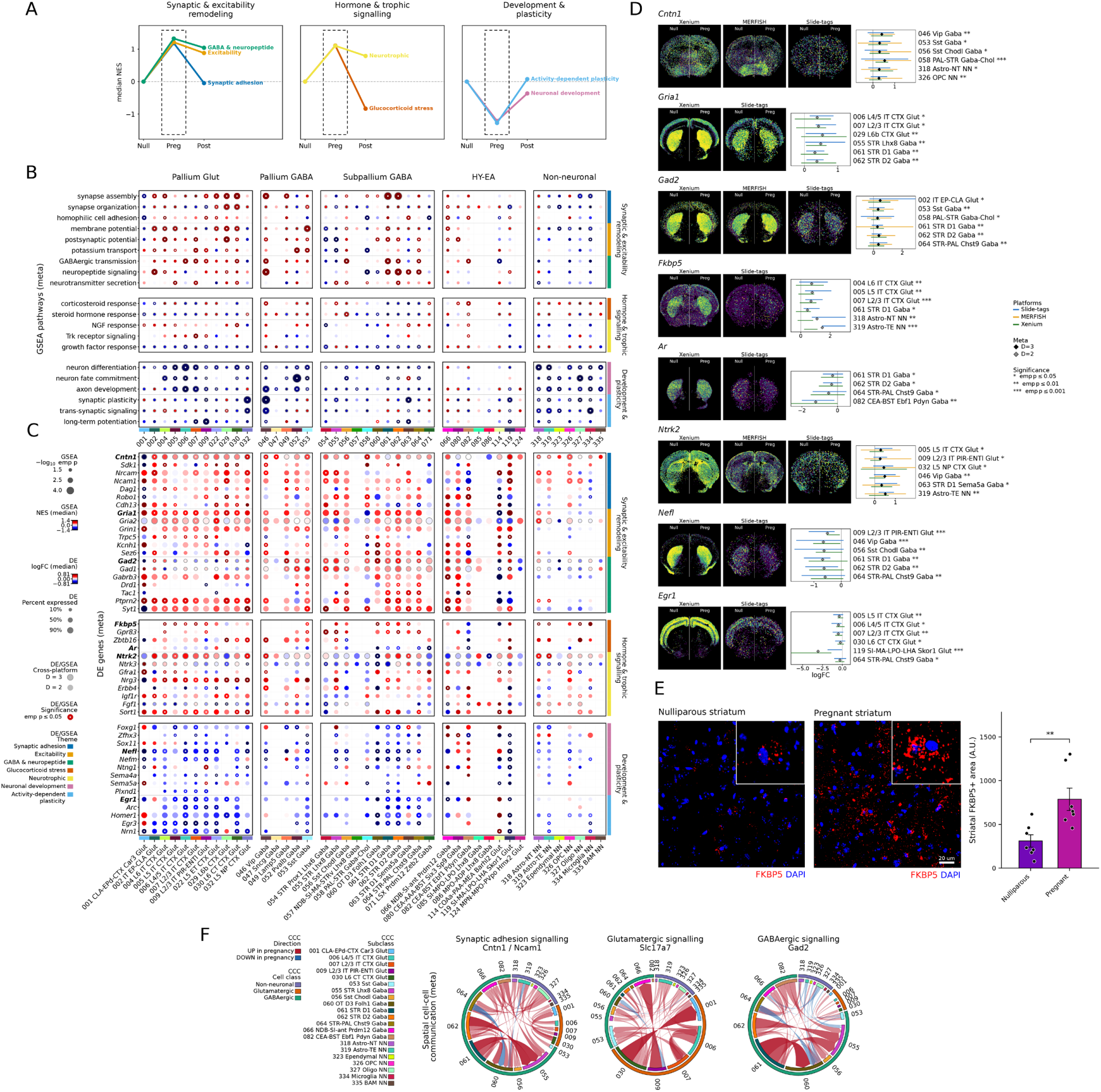
Late pregnancy remodels synaptic, hormonal and developmental programs in maternal neurons. **A)** Peripartum trajectories of neuronal programs. Each line traces one GO biological-process theme’s median normalized enrichment score (NES) across the neuronal cell types enriched for it in pregnancy, at three reproductive states (nulliparous, Null; pregnant, Preg; postpartum, Post) referenced to the nulliparous baseline (NES = 0); themes are grouped into the super-groups labeled above each panel. Programs that increased in pregnancy remained elevated after birth (excitability, GABA and neuropeptide, neurotrophic), returned toward baseline (synaptic adhesion) or decreased below it (glucocorticoid stress); programs decreased in pregnancy recovered postpartum (neuronal development, activity-dependent plasticity). The dashed box marks the pregnant state detailed in B–F (postpartum profiled on MERFISH and Slide-tags only). **B)** Gene set enrichment (GSEA) for pregnant versus nulliparous across GO biological-process terms, grouped into the same super-groups (right). Columns are the neuronal and glial subclasses with significant enrichment, grouped by brain region. Dot size, significance (−log₁₀ empirical *p*); fill color, median NES (red, increased; blue, decreased in pregnancy); ring, reproduced on all three platforms; white center dot, *p* ≤ 0.05. **C)** Differential expression of genes spanning the same super-groups; those in D are bold. Dot size, fraction of cells expressing the gene; fill color, median log₂ fold change; ring and center dot as in B. **D)** Spatial expression of representative genes. Single-cell expression in nulliparous (left of midline) and pregnant (right) hemispheres for every platform assaying it, beside per-platform and meta-analyzed log₂ fold changes (95% CI) across the cell types in which it is significant. **E)** Immunofluorescence validation of striatal FKBP5 (red) with DAPI (blue) in nulliparous and pregnant animals (inset, magnified cell; scale bar, 20 µm), with FKBP5⁺ area quantified (arbitrary units; bars, mean ± SEM; points, animals; **, *p* ≤ 0.01, unpaired t-test). **F)** Spatial cell–cell communication. Chord diagrams for synaptic adhesion (*Cntn1*/*Ncam1*), glutamatergic (*Slc17a7*) and GABAergic (*Gad2*) signaling show ligand–receptor interactions changing consistently in pregnancy across Slide-tags and Xenium among the most active neuronal subclasses and their glial partners; ribbon color, direction (red, increased; blue, decreased); width, magnitude; arrowheads point from sender to receiver.

Genes for synapse assembly increased expression in cortical and striatal neurons (synaptic-adhesion theme increased in 18 of the 42 neuronal subclasses, median normalized enrichment score (NES) +1.23, and in no non-neuronal subclass; Fig. 3B). One such example was *Cntn1*, an adhesion molecule that builds and stabilizes synapses^33^, which increased expression in many cell types on all three platforms (Fig. 3D). Receptors and enzymes for fast transmission also increased expression, including the AMPA-receptor subunit *Gria1* and the GABA-A receptor subunit *Gabrb3* across the cortex and striatum, and the GABA-synthesizing enzyme *Gad2* in striatal and cortical inhibitory neurons, echoing the synaptic protein gains reported in the maternal rodent brain^34^. Cell-cell signaling corroborated these changes: adhesion, glutamatergic and GABAergic signals all increased, with adhesion and GABAergic signaling converging on striatal *Lhx8* neurons (Fig. 3F; Supplementary Data 5). We then tracked each theme across reproductive states as its median NES over the neuronal subclasses enriched for it in pregnancy, relative to the nulliparous level (Fig. 3A). The synapse-assembly theme returned to that level after birth (+1.18 in pregnancy, −0.05 postpartum), whereas the excitability and GABAergic themes persisted (+1.21 to +0.88 and +1.32 to +1.04).

Pregnancy also increased corticosteroid response in 5 of the 42 neuronal subclasses and steroid hormone response in 3 (median NES +1.16 and +1.04; Fig. 3B). The glucocorticoid-response genes *Fkbp5*, *Gpr83* and *Zbtb16* increased expression across several cell types (Fig. 3C,D). *Fkbp5* encodes a co-chaperone that reduces the affinity of the glucocorticoid receptor for cortisol and is itself induced by receptor activation, a negative feedback loop. Variants that increase its induction are associated with affective and anxiety disorders^35^. Immunostaining confirmed the increase at the protein level, with FKBP5 increasing about 2.5-fold in the striatum (*p* = 0.0092, Welch t-test; Fig. 3E). In the same striatal and amygdalar neurons, expression of the androgen receptor *Ar* decreased (Fig. 3C), suggesting a rebalancing of steroid tone at stress- and social-behavior circuits^36^. Neurotrophic signaling was reconfigured in parallel, though the theme moved in both directions across subclasses (increased in 7 neuronal subclasses, median NES +1.17; decreased in 6, median NES −1.12): neurons demonstrated increased expression of the receptors for BDNF, GDNF and IGF-1 (*Ntrk2*, *Gfra1*, *Igf1r*), while the neurotrophin-3/TrkC receptor *Ntrk3* decreased in neurons and glia alike (Fig. 3C). In the postpartum period, the two responses diverged: the glucocorticoid program reversed below the nulliparous level while the neurotrophic shift persisted (Fig. 3A).

The preoptic area and its neighboring nuclei house the circuits that drive parental care^11^. We captured 19 of these subclasses across the preoptic area, extended amygdala, lateral septum, basal forebrain and olfactory tubercle, each identified on all three platforms (Supplementary Fig. 6A,B). Their defining markers were strongly expressed in the subclasses where they were detected but unchanged by pregnancy: galanin (*Gal*), the calcitonin receptor (*Calcr*), the progesterone receptor (*Pgr*) and neurokinin B (*Tac2*). Across the six preoptic subclasses the number of reproducible changes varied considerably, from 237 genes in 119 SI-MA-LPO-LHA Skor1 Glut to 3 in 118 ADP-MPO Trp73 Glut (Supplementary Data 3). In pregnant relative to nulliparous mice, the neuropeptides *Nts* and *Crh* increased across multiple subclasses. Two hormone receptors also changed: *Ar* decreased in the lateral septum and extended amygdala, and the thyroid-hormone receptor *Thrb* increased in the lateral septum and preoptic area (Supplementary Fig. 6C). Changes in the remaining steroid and peptide receptors were sparse: *Esr1* and *Oxtr* each decreased in a single subclass and *Prlr* did not change. Beyond these effectors, the maternal subclasses participated in the programs described above, with adhesion, postsynaptic transmission and ion transport recurring among their enriched terms (Supplementary Data 4). These circuits therefore retain their defining markers through pregnancy while changing along with the rest of the tissue.

In pregnant relative to nulliparous mice, neurons decreased expression of two other sets of genes. Neuronal-development terms decreased in 22 of the 42 neuronal subclasses and in 10 of the 11 non-neuronal subclasses (median NES −1.21), and activity-dependent-plasticity terms in 9 neuronal and 9 non-neuronal subclasses (median NES −1.32); these were the most widely shared changes we detected. Developmental genes related to axonal growth and cytoskeletal assembly decreased across most cell types (*Nefl*, *Nefm*, *Gap43* and *Tubb3*; Fig. 3C,D; Supplementary Data 3), a sign of reduced structural growth in these already-mature neurons^37^. Genes associated with activity-dependent signaling (*Egr1*, *Egr3*, *Arc* and *Fos*) and genes supporting long-term potentiation (*Homer1* and *Nptx2*) also decreased expression (Fig. 3C,D). Astrocytes, oligodendrocytes and microglia decreased the same two sets of pathways, including the axon-guidance and synapse-shaping genes through which they support neuronal circuits (Fig. 3B). These decreases occurred in the same neurons that were changing in synapse-related gene expression. Striatal D1 medium spiny neurons, for example, increased *Gria1* while decreasing *Egr1* and *Nefl* expression (Fig. 3C). The two sets diverged in the postpartum: immediate-early and plasticity genes recovered to the nulliparous level, whereas the decrease in axon-growth and cytoskeletal genes only partly recovered (median NES −1.21 in pregnancy and −0.36 postpartum; Fig. 3A). The neuronal program therefore has two layers. One is transient, peaking in pregnancy and then resetting postpartum, including synapse assembly, the glucocorticoid response and the suppression of immediate-early and plasticity genes (Fig. 3A). The other layer is consolidated and carries into motherhood, including expression changes suggestive of stronger excitability and inhibition, sustained neurotrophic support and a decrease in axon-growth and cytoskeletal genes that only partly recovers.

### An immune and vascular signaling program alters cell state without remodeling the vascular bed

In pregnancy, non-neuronal cells upregulated both an immune and a vascular program (innate immune activation increased in all 11 non-neuronal subclasses, median NES +1.32; cytokine signaling in 10 and myeloid effector programs in 10, median NES +1.29 and +1.21) (Fig. 4B; Supplementary Data 4).

**Fig. 4.**
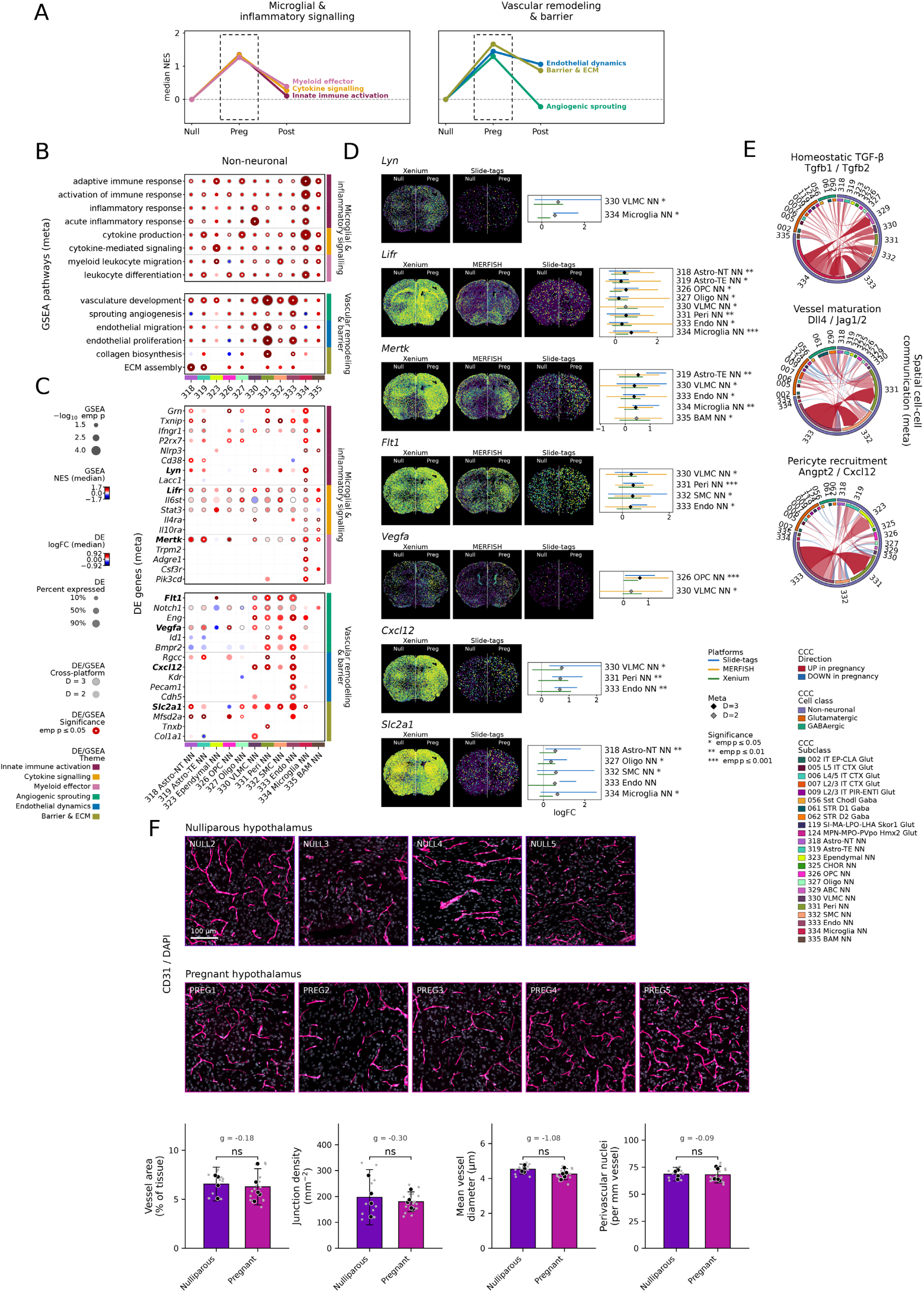
**Late pregnancy activates inflammatory and angiogenic programs in maternal glia and vasculature without altering vessel architecture. *A)*** Peripartum trajectories of glial–vascular programs, plotted as in Fig. 3A across the non-neuronal cell types enriched for each theme in pregnancy; super-groups are labeled above each panel. All programs increased in pregnancy; the microglial themes (innate immune activation, cytokine signaling, myeloid effector) and angiogenic sprouting then returned toward baseline, whereas endothelial dynamics and barrier–ECM remained elevated postpartum. The dashed box marks the pregnant state detailed in B–F (postpartum profiled on MERFISH and Slide-tags only). **B)** Gene set enrichment (GSEA) for pregnant versus nulliparous across GO biological-process terms, grouped into the same super-groups (right). Columns are the eleven non-neuronal subclasses of the glial and vascular niche. Dot size, significance (−log₁₀ empirical p); fill color, median NES (red, increased; blue, decreased in pregnancy); ring, reproduced on all three platforms; white center dot, p ≤ 0.05. **C)** Differential expression of genes spanning the same super-groups, with those in D in bold; encoding as in Fig. 3C. **D)** Spatial expression of representative genes in nulliparous (left of midline) and pregnant (right) hemispheres for every platform assaying it, beside per-platform and meta-analyzed log₂ fold changes (95% CI) across the cell types in which the gene is significant. **E)** Spatial cell–cell communication. Chord diagrams for homeostatic TGF-β (Tgfb1/Tgfb2), vessel maturation (Dll4/Jag1/Jag2) and pericyte recruitment (Angpt2/Cxcl12) show ligand–receptor interactions changing consistently in pregnancy across Slide-tags and Xenium among the most active glial and vascular subclasses and their neuronal partners; encoding as in Fig. 3F. **F)** Immunofluorescence validation of vascular architecture in the preoptic area. CD31 (magenta) and DAPI (gray) in a representative field from an independent cohort of four nulliparous and five pregnant mice (scale bar, 100 µm), with vessel area, junction density, mean diameter and perivascular nuclei per vessel length quantified (bars, mean ± 95% CI; black points, animal means; gray points, the 28 fields; g, Hedges’ g; ns, not significant, unpaired t-test on animal means). No measure differed between states, here or across all eighteen measures (Supplementary Fig. 7; Supplementary Data 7).

The immune-signaling genes that increased in microglia included receptors for inflammatory and anti-inflammatory cytokines (*Ifngr1*, *Il4ra*, *Il10ra*), signaling kinases (*Lyn*, *Pik3cd*), inflammasome components (*P2rx7*, *Trpm2*, *Nlrp3*) and machinery for lysosomal degradation and non-inflammatory clearance of dying cells (*Grn*, *Mertk*). *Lifr*, which encodes the receptor for leukemia inhibitory factor, was upregulated in nearly every non-neuronal cell type, as were the genes for its partner subunit gp130 (*Il6st*) and the transcription factor STAT3 (*Stat3*), spanning successive components of the JAK-STAT pathway (Fig. 4C,D). TGF-β signaling from endothelial cells and astrocytes to microglia increased during pregnancy (Fig. 4E). TGF-β is required for the molecular signature that distinguishes microglia from other myeloid cells^38^. These changes occurred without a detectable shift in microglial abundance, suggesting an altered immune state within a stable population rather than microglial expansion (Supplementary Data 2).

In pregnant relative to nulliparous mice, endothelial cells, mural cells (pericytes and smooth muscle) and vascular leptomeningeal cells (VLMCs) increased expression of genes associated with angiogenesis and barrier function (angiogenic sprouting increased in 7 of the 11 non-neuronal subclasses, endothelial dynamics in 4 and barrier and extracellular-matrix programs in 4; median NES +1.27, +1.45 and +1.59) (Fig. 4B). In endothelial cells and pericytes, expression of the VEGF receptors *Flt1* and *Kdr* increased, along with the TGF-β co-receptor endoglin (*Eng*) and *Rgcc*, a regulator of endothelial proliferation (Fig. 4C,D). Expression of *Vegfa*, which encodes the ligand for these receptors, increased in oligodendrocyte precursors and VLMCs. Endothelial cells increased their signaling to mural cells through the Notch ligand *Dll4*, and to pericytes and ependymal cells through the chemokine *Cxcl12* (Fig. 4E; Supplementary Data 5). Pericytes and VLMCs also increased matrix components of the vessel wall (*Tnxb*, *Col1a1*). The cells carrying this program did not change in number or position: vascular cell type abundance and neighborhood composition were unchanged (Supplementary Fig. 5; Supplementary Data 2). In development, these are the pathways that govern mural-cell recruitment and vessel maturation^39^, so we asked whether the vascular bed itself had changed.

To test whether this signaling program remodels the vasculature itself, we immunostained the preoptic area for the endothelial marker PECAM-1 (CD31) in an independent cohort of nulliparous and late-pregnant mice (embryonic day 17.5 to 18.5; 4 and 5 animals, 28 fields; Fig. 4F). However, we detected no statistically significant difference in the structure of the vascular bed. Vessels occupied 6.6% of the tissue in nulliparous mice and 6.3% in pregnant mice (Welch t-test on animal means, *p* = 0.76) and perivascular nuclei per unit vessel length were likewise unchanged (68.6 and 68.1 per mm, *p* = 0.88). None of 18 morphometric measures, spanning vessel density, network architecture, caliber, tissue supply and cellularity, differed after correction for multiple testing (Supplementary Fig. 7; Supplementary Data 7). Transcriptional state and vascular architecture therefore moved independently, a dissociation we consider further in the Discussion.

After birth the endothelial and barrier programs remained elevated (+1.45 to +1.06 and +1.67 to +0.87), whereas the angiogenic program reversed below the nulliparous level (+1.30 to −0.22) and the immune programs returned toward it (postpartum medians +0.11 to +0.39; Fig. 4A). Pregnancy therefore changes the signaling state of vascular and immune cells without changing their numbers or the structure of the vascular bed.

### A sustained lipid program increases lipid turnover and shifts cholesterol from synthesis to uptake

Pregnancy reprogrammed lipid metabolism along two axes, spanning both neurons and glia (Fig. 5B; Supplementary Data 4). Along the first, intrinsic lipid metabolism increased through coordinated changes in the enzymes that build and turn over brain lipids (membrane-lipid metabolism increased in 12 neuronal and 3 non-neuronal subclasses, sphingolipid metabolism in 11 and 2, and fatty-acid catabolism in 7 and 4; median NES +1.20, +1.24 and +1.36; Fig. 5B), consistent with the drawdown of maternal brain lipid across pregnancy and lactation^40^. Membrane-lipid metabolism showed the largest increase, led by *Mfsd2a*, which transports docosahexaenoic acid into the brain for membrane synthesis^41^, in astrocytes and oligodendrocytes (Fig. 5C,D). The glycerolipid-synthesizing enzyme *Lpin1* increased alongside it in astrocytes, cortical excitatory neurons and striatal neurons (Fig. 5C). Sphingolipid turnover increased in parallel, marked by the lysosomal galactosidase *Glb1* and the ganglioside-catabolizing hexosaminidase *Hexa*, in keeping with the specialized sphingolipid composition of glial and myelin membranes^42^ (Fig. 5C). Fatty-acid catabolism increased most in glia, led by the branched-chain and fatty-acid catabolic enzyme *Ivd* in OPCs, astrocytes and pericytes (Fig. 5C). One gene ran counter: *Fabp7*, which binds and shuttles fatty acids rather than degrading them, decreased all 10 subclasses in which it changed and most strongly in astrocytes (Fig. 5C).

**Fig. 5.**
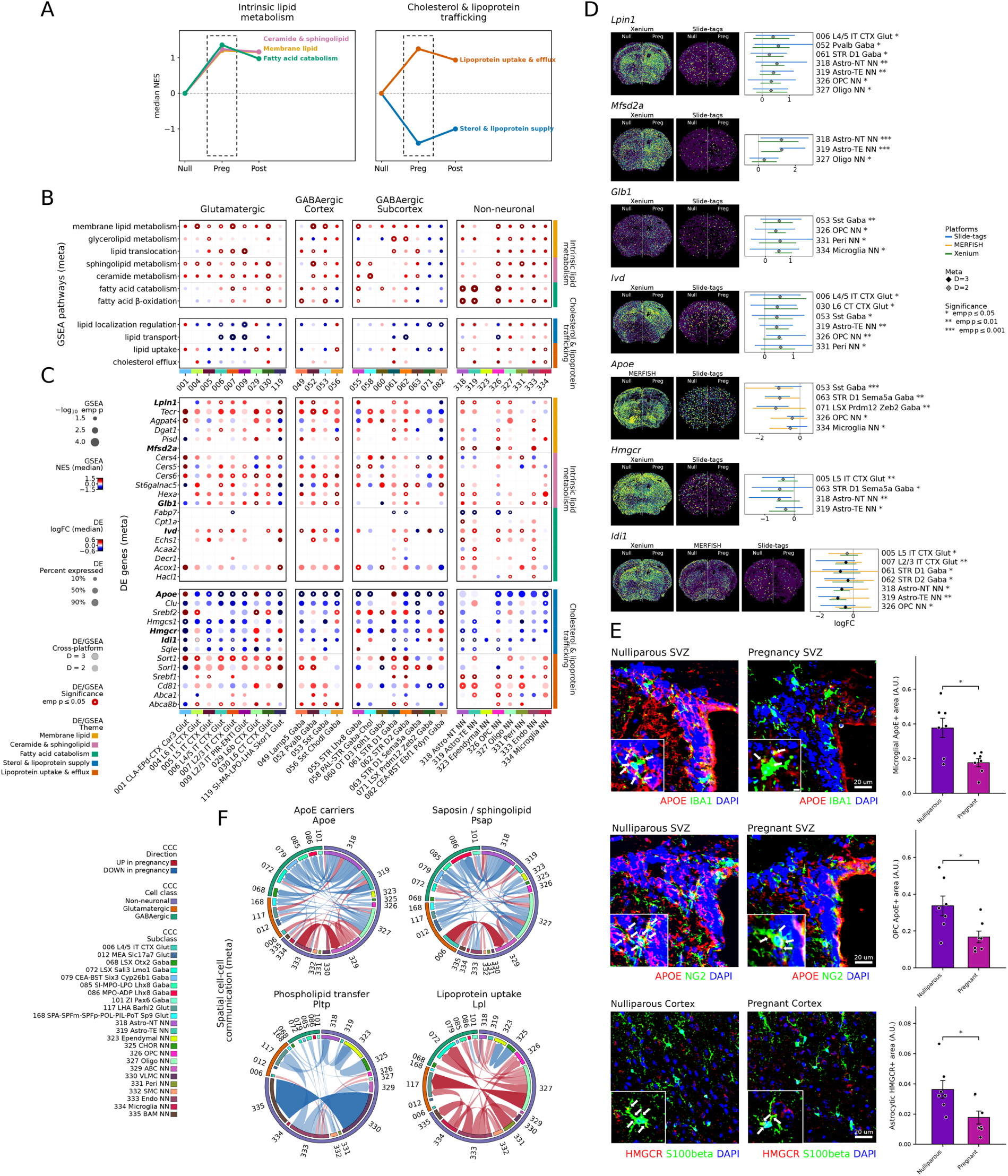
**Late pregnancy reprograms lipid metabolism in neurons and glia, suppressing cholesterol supply while elevating lipid programs that largely persist postpartum. *A)*** Peripartum trajectories of the lipid-metabolism programs, plotted as in Fig. 3A across the cell types enriched for each theme in pregnancy; the two super-groups (intrinsic lipid metabolism; cholesterol and lipoprotein trafficking) are labeled above each panel. Cholesterol synthesis and supply decreased in pregnancy, whereas the membrane-lipid, sphingolipid, fatty-acid and lipoprotein-uptake themes increased, and most remained shifted postpartum. The dashed box marks the pregnant state detailed in B–F (postpartum profiled on MERFISH and Slide-tags only). **B)** Gene set enrichment (GSEA) for pregnant versus nulliparous across GO biological-process terms, grouped into the same super-groups (right). Columns are the neuronal and glial subclasses with significant enrichment, grouped by cell class (glutamatergic, GABAergic cortical, GABAergic subcortical, non-neuronal). Dot size, significance (−log₁₀ empirical *p*); fill color, median NES (red, increased; blue, decreased in pregnancy); ring, reproduced on all three platforms; white center dot, *p* ≤ 0.05. **C)** Differential expression of genes spanning the same super-groups, with those in D in bold; encoding as in Fig. 3C. **D)** Spatial expression of representative genes in nulliparous (left of midline) and pregnant (right) hemispheres for every platform assaying it, beside per-platform and meta-analyzed log₂ fold changes (95% CI) across the cell types in which the gene is significant. **E)** Immunofluorescence validation of the glial cholesterol-supply program in nulliparous and pregnant animals: APOE (red) in IBA1⁺ microglia and in NG2⁺ oligodendrocyte precursors (green) in the subventricular zone, and HMGCR (red) in S100β⁺ astrocytes (green) in the cortex, with DAPI (blue) counterstain. Insets, magnified cells with arrows; scale bars, 20 µm; bars, mean ± SEM; points, animals; *, *p* ≤ 0.05, unpaired t-test (Supplementary Data 6). **F)** Spatial cell–cell communication. Chord diagrams for APOE carriers (*Apoe*), saposin and sphingolipid signaling (*Psap*), phospholipid transfer (*Pltp*) and lipoprotein uptake (*Lpl*) show ligand–receptor interactions changing consistently in pregnancy across Slide-tags and Xenium among the most active glial subclasses and their neuronal partners; encoding as in Fig. 3F.

Along the second axis, cholesterol synthesis and supply decreased while lipoprotein uptake increased (sterol and lipoprotein supply decreased in 6 neuronal and 1 non-neuronal subclass, median NES −1.40, while lipoprotein uptake and efflux increased in 2 neuronal and 3 non-neuronal subclasses, median NES +1.25; Fig. 5B). The maternal brain normally synthesizes its own cholesterol locally, behind the blood-brain barrier^43^, and astrocytes deliver it to neurons on APOE-containing lipoprotein particles to support synapse formation^44^. *Apoe*, the principal lipid and cholesterol carrier of the brain, decreased in 22 subclasses and increased in none, making it the most pervasive lipid-related change we detected across cortical excitatory neurons, interneurons, striatal neurons, microglia and OPCs (Fig. 5C). The cholesterol-synthesis pathway decreased with it, marked by the sterol-synthesis isomerase *Idi1* and the rate-limiting enzyme *Hmgcr* in deep-layer cortical neurons, astrocytes and striatal neurons (Fig. 5C). The receptors and transporters that import and redistribute lipoprotein increased instead, including the sorting receptors *Sort1* and *Sorl1* and the cholesterol-efflux transporter *Abca1* in oligodendrocytes, astrocytes and OPCs (Fig. 5C). We confirmed the decrease at the protein level: APOE decreased by 53% in microglia and 50% in OPCs, and the rate-limiting enzyme HMGCR by 51% in astrocytes (Welch t-test, *p* = 0.010, 0.022 and 0.027; n = 7 mice per group; Fig. 5E; Supplementary Data 6).

Cell-cell communication, inferred from the expression of each ligand in sending cells and of its receptors in receiving cells, changed in the same directions (Fig. 5F; Supplementary Data 5). Signaling from lipoprotein lipase (*Lpl*) to its receptors *Lrp1* and *Vldlr* increased in pregnancy. *Lpl* mediates lipoprotein uptake in the central nervous system^45^. By contrast, signaling from *Apoe* to its receptors *Lrp1* and *Sorl1*, and from prosaposin (*Psap*) to *Sort1* and *Gpr37l1*, decreased. This signaling flowed predominantly from glia to neurons, matching the same division of labor: glia carried lipoprotein uptake and neurons led the decrease in cholesterol supply. Both axes were established in pregnancy and retained most of their magnitude into the postpartum period (Fig. 5A). Together, these results suggest a coordinated, pan-cellular reprogramming of brain lipid metabolism that increases membrane-lipid turnover, shifts cholesterol provisioning from synthesis to uptake, and outlasts pregnancy itself.

### Genetic risk for psychiatric disorders concentrates in neurons and shifts across reproductive states

Depression is common in pregnancy and the postpartum period, including in mothers with no prior history of the disorder^9,10^. To identify the cells that carry the genetic risk for depression, we applied gsMap to the Slide-tags and Xenium datasets, integrating the spatial transcriptome with GWAS summary statistics for 16 human neurological and psychiatric traits across the three reproductive states^46^ (Supplementary Data 8). Major depressive disorder (MDD), neuroticism, ADHD and autism showed the strongest trait enrichment (Fig. 6A). A GWAS for Postpartum depression was not enriched, however we reason that this may be due to the much smaller sample sizes of this GWAS compared to others (48,000 individuals against 500,200 for MDD).

**Fig. 6.**
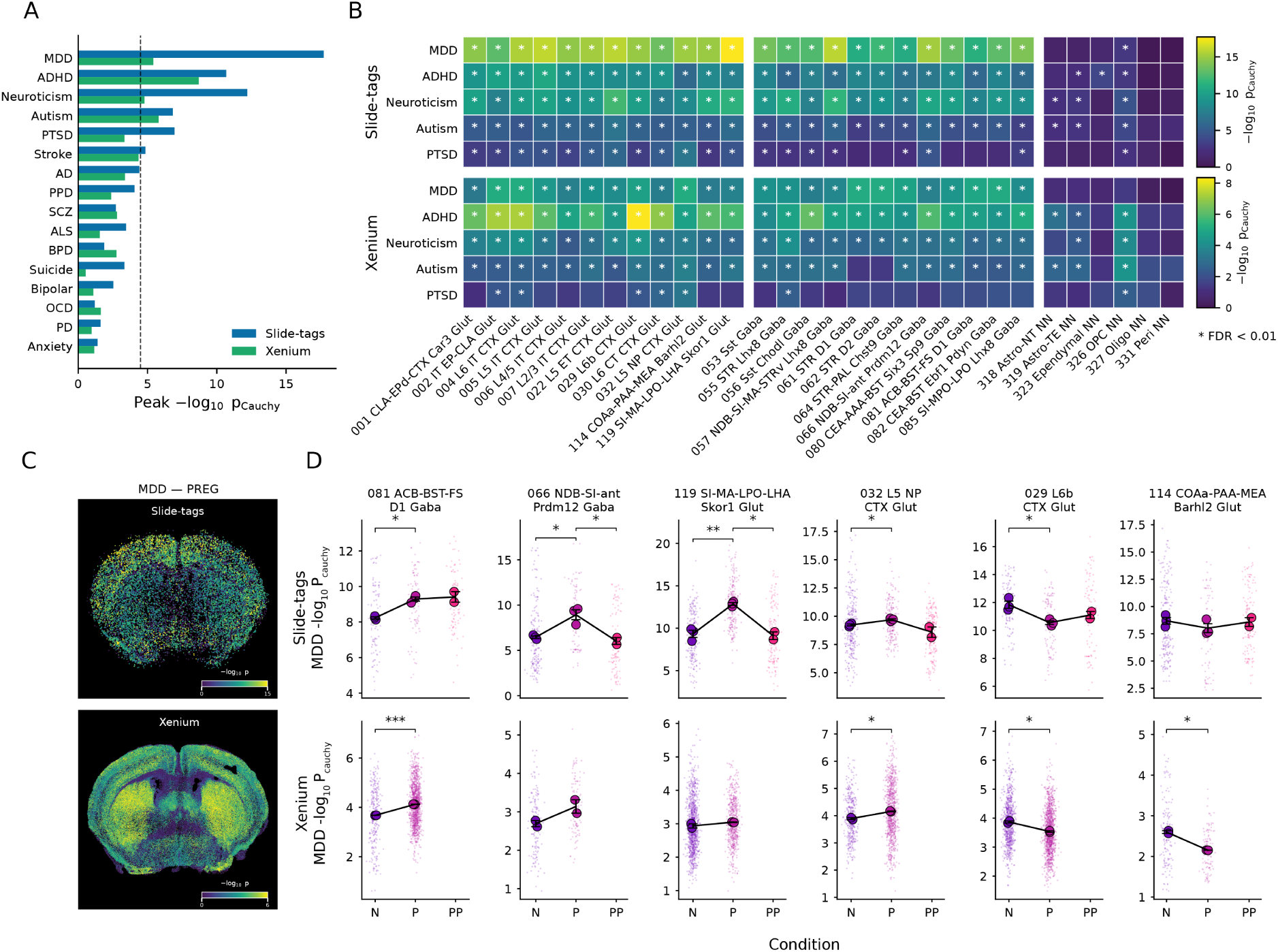
Spatially resolved mapping of disease risk across cell types and reproductive states. GWAS summary statistics for 16 human neurological and psychiatric traits were integrated with the Slide-tags and Xenium spatial transcriptomes using gsMap^46^. **A)** Peak association per trait, the maximum −log₁₀ *p* Cauchy across all cell subclasses and reproductive states, for Slide-tags (blue) and Xenium (green). Dashed line, Bonferroni threshold (α = 0.05) across all traits and subclasses tested. **B)** Trait × cell type associations for the five leading traits in Slide-tags (top) and Xenium (bottom). Fill color, maximum −log₁₀ *p* Cauchy across reproductive states, scaled independently per platform; asterisks, FDR < 0.01. **C)** Per-cell MDD association in the pregnant state, plotted at registered coordinates, for Slide-tags (top) and Xenium (bottom). **D)** MDD association across reproductive states for six exemplar subclasses, in Slide-tags (top) and Xenium (bottom). Small points, individual cells; large points, per-sample means; line and error bars, mean ± SEM of sample means. Nulliparous (N), pregnant (P), postpartum (PP); Xenium did not sample the postpartum state. Brackets, per-platform comparisons (* *p* < 0.05; ** *p* < 0.01; *** *p* < 0.001).

Across these traits, genetic risk was carried almost entirely by neurons (Fig. 6B). For MDD, the Cauchy-combined association reached FDR <0.01 in 55 of 56 neuronal subclasses on Slide-tags and 78 of 82 on Xenium, but in 1 of 12 and none of 16 non-neuronal subclasses. The strongest associations implicated cortical excitatory neurons from layer 2/3 through layer 6, GABAergic neurons of the basal forebrain (NDB-SI-MA-STRv Lhx8 Gaba, NDB-SI-ant Prdm12 Gaba), and excitatory neurons of the substantia innominata and lateral preoptic area (SI-MA-LPO-LHA Skor1 Glut), the most enriched subclass overall (Fig. 6B,C). A recent trans-ancestry depression GWAS reports the same neuronal and amygdalar enrichment^47^.

We next asked whether the MDD associations changed across reproductive states — that is, whether pregnancy altered which cells most distinctively express the genes carrying depression heritability (Fig. 6D). In pregnancy the association strengthened in four subclasses: D1-expressing GABAergic neurons of the nucleus accumbens and bed nucleus of the stria terminalis (ACB-BST-FS D1 Gaba), layer 5 near-projecting cortical neurons (L5 NP CTX Glut), GABAergic neurons of the diagonal band and substantia innominata (NDB-SI-ant Prdm12 Gaba), and excitatory neurons of the substantia innominata and lateral preoptic area (SI-MA-LPO-LHA Skor1 Glut). It weakened in two other subclasses, one cortical (L6b CTX Glut) and one amygdalar (COAa-PAA-MEA Barhl2 Glut). Slide-tags and Xenium agreed on the direction of change in all six subclasses. In Slide-tags, the change in the basal forebrain and preoptic subclasses was transient, returning toward the nulliparous level in the late-postpartum period. The subclasses whose association strengthened during pregnancy sit in the nucleus accumbens and bed nucleus of the stria terminalis, which govern motivation and the stress response^48,49^, and in the basal forebrain and preoptic area, which control maternal behavior^12^. Together, these results place the genetic risk for depression and related disorders in neurons distributed across the cortex, basal forebrain, striatum and amygdala of the maternal brain, and not in glial or vascular cells.

## Discussion

Our study provides a cell type-resolved view of the maternal mouse brain across the peripartum transition, built from three spatial technologies annotated under a common atlas and combined by cross-platform meta-analysis. Neurons increased expression of genes for synapse assembly and shifted toward stronger excitability and inhibition while suppressing genes for activity-dependent plasticity and structural growth. Glia and vascular cells engaged a coordinated immune and angiogenic signaling program without remodeling the vascular bed itself. Neurons and glia together shifted cholesterol provisioning from local synthesis to lipoprotein uptake while increasing membrane-lipid turnover. These programs differed in persistence: some, including synapse assembly and the glucocorticoid response, reset after birth, whereas others, including the shifts in excitability, endothelial and barrier signaling, and lipid handling, carried into lactation. Mapping human depression genetics onto these data placed genetic risk almost entirely in neurons, and this risk shifted with reproductive state in circuits, including the preoptic area, basal forebrain, and nucleus accumbens and bed nucleus of the stria terminalis, that are simultaneously being remodeled for maternal care.

Cellular composition, local arrangement and vascular architecture were each tested directly, and none changed appreciably in the regions we sampled. However, structural remodeling of the maternal brain need not alter which cells occupy the tissue. Perineuronal nets assemble in the preoptic area through gestation and fade after birth^50^, and hippocampal dendritic spine density increases in late pregnancy and lactation^51^. Compositional change does occur in the neurogenic niches, which lie outside our coronal plane, where prolactin drives progenitor production^52^ and a recent single-cell survey of the female mouse hippocampus finds its clearest compositional changes among dentate gyrus stem cells^16^. Pregnancy therefore appears to remodel the maternal brain from within its resident cells rather than by altering their number or arrangement.

In pregnant relative to nulliparous mice, neurons engaged programs for synapse assembly while suppressing those for activity-dependent plasticity and structural growth. Pregnancy hormones produce both halves of this pattern. Estradiol replacement in ovariectomized rodents increases synapse number in the hippocampus of adult females^53^. In the preoptic area, pregnancy hormones promote dendritic spine formation and excitatory synaptic input while sparsening population activity in vivo^13^. Many of the cell types engaging the synaptic program also mounted a coordinated glucocorticoid response, matching the increase in basal corticosterone that accompanies late pregnancy^54^. The synaptic and glucocorticoid responses did not persist into late lactation, whereas the changes in excitability, inhibition and structural growth did. The pregnant animals had not yet encountered pups, so this remodeling precedes the behavior it may support.

In pregnancy, expression of vascular endothelial growth factor (VEGF) receptors increased across the neurovascular unit. Immunostaining of the preoptic area found no change in vessel density, caliber, network architecture or perivascular cellularity. The vascular compartment therefore changed its transcriptional state without changing its structure. In the rat, by contrast, pregnancy causes outward remodeling of penetrating arterioles and higher capillary density^55^. *Flt1* was the most consistently upregulated of the VEGF receptors, reproducing on every platform in each vascular type. It encodes both the membrane receptor and the soluble decoy whose circulating form blocks VEGF-induced permeability at the maternal cerebral barrier^56^. That barrier fails in eclampsia, a leading cause of maternal death in which the cerebral circulation is the primary effector^57^. Determining which isoform dominates would show whether the maternal cerebral vasculature is more responsive to VEGF or better protected from it.

Pregnancy and lactation can deplete the maternal brain of docosahexaenoic acid, a principal constituent of neuronal membranes^40^. In pregnant relative to nulliparous mice, the enzymes that build membrane lipids and those that degrade them increased together in neurons and glia, suggesting higher turnover. Cholesterol is another constituent of those membranes, and its synthesis decreased while the receptors that take up lipoprotein particles increased. Unlike docosahexaenoic acid, which enters from the circulation^41^, almost all brain cholesterol is made locally and recycled between cells on apolipoproteins^43^. A shift toward uptake therefore redistributes the existing pool rather than replenishing it. Neurons make enough cholesterol to survive but not enough to build many synapses, and glia supply the shortfall on APOE-containing lipoproteins^44^. *Apoe* was the most pervasive lipid change we detected, and it decreased while neurons were assembling synapses. Astrocytes held *Apoe* steady and decreased cholesterol synthesis instead. Glia upregulated the transporter that loads cholesterol onto lipoprotein, whereas neurons upregulated the receptors that take it up. Immunostaining confirmed both decreases where the transcripts changed, APOE in microglia and oligodendrocyte precursors and HMGCR, the rate-limiting enzyme of cholesterol synthesis, in astrocytes. Both persisted into late lactation, when milk export continues to draw on maternal lipid. Neurons may have raised these receptors because fewer particles were available rather than because more cholesterol was moving between cells. Isotope labeling of cholesterol transfer between glia and neurons would distinguish the two.

To connect these programs to human disease, we mapped depression genetics onto the mouse cell types^46^. Genes carrying depression heritability were most distinctively expressed in neurons of the cortex, striatum, basal forebrain and amygdala, and scarcely at all in glial or vascular cells. A trans-ancestry study of depression implicates the same neuronal classes, including amygdalar neurons, in both mouse and human data^47^. The mapping identifies where those genes are concentrated, not a change in risk itself. In six subclasses the strength of that association changed with reproductive state, and both platforms agreed on the direction. They include populations of the preoptic area and basal forebrain, which control maternal behavior^11^, and the nucleus accumbens and bed nucleus of the stria terminalis, which govern motivation and the stress response^48,49^. The preoptic population most strongly associated with depression risk is excitatory, and preoptic glutamatergic neurons regulate sleep and body temperature as well as parental care^58,59^. Pregnancy may therefore concentrate depression-associated expression in circuits it is simultaneously remodeling for maternal care.

This study has several limitations. Each animal contributed a single coronal plane, so the changes we describe are limited to the structures it crosses, although registration showed the sampled planes to be closely matched between conditions. Our significance threshold is exploratory, so the genes we name are candidates supported by cross-platform reproducibility and pathway enrichment. Our resolution is the subclass, so changes confined to hormone-receptor-defined subsets within a subclass may be missed, and enriching for such neurons recovers state-dependent expression that unenriched profiling of these nuclei does not^60^. Xenium did not sample the postpartum state, so persistence rests on a smaller gene space and fewer animals. Nulliparous animals were staged to diestrus, but lactation suppresses the estrous cycle, so our postpartum animals were in anestrus and that comparison reflects ovarian steroid state as well as parity. Sampling in the weeks after delivery, when psychiatric admission is most frequent^8^, would cover the window of highest human risk. The genetic mapping also crosses species that differ in reproductive physiology. Serial sampling, whole-brain profiling and functional manipulation will be needed to connect these programs to maternal behavior.

The maternal brain we describe is reconfigured rather than rebuilt, its cells changing state while their number and arrangement hold. The neurons that carry depression risk sit among the circuits being remodeled, so adaptation and vulnerability may occupy the same cells rather than merely the same window. Identifying those cells is a step toward explaining the psychiatric risk of the peripartum period.

## Methods

### Mouse husbandry and ethics

All animal procedures were approved by the SickKids Research Institute Animal Care Committee. Female C57BL/6J mice (Jackson Laboratories) were acclimated for one week under a 12:12 light-dark cycle with ad libitum food and water. Brain tissue was collected at three maternal stages: nulliparous (virgin), timed pregnant (gestational day 18) and postpartum (day 20, pre-weaning). For euthanasia, virgin females were approximately 9-10 weeks of age; pregnant mice were approximately 9 weeks of age; post-partum mice were approximately 11 weeks of age. Virgin mice were group-housed without males to suppress and synchronize the estrous cycle, exposed to soiled male bedding for 72 h to induce synchronized estrus (Whitten effect), monitored daily by vaginal cytology (lavage, wet-mount smear, cell morphology) and euthanized on confirmed entry into diestrus. Pregnant and postpartum females were single housed from gestational day 10 until euthanasia, and the stud male was removed at plug detection. Timed breeding occurred in house. Litter sizes ranged from 5-8 offspring per dam. We profiled three mice per condition by MERFISH (n = 9); three virgin, three pregnant and two postpartum mice by Slide-tags (n = 8, one postpartum brain being unavailable for processing); and two nulliparous and two pregnant mice by Xenium (n = 4, sampling those two states only) (Supplementary Data 1). Brains were embedded in optimal cutting temperature compound (OCT) immediately after euthanasia, flash-frozen on dry ice and stored at −80 °C until sectioning.

### Slide-tags tissue and library preparation

Fresh-frozen tissue was coronally sectioned at 25-30 µm in an RNase-free cryostat (Leica CM1850) at −18 °C onto pre-chilled 10x10 mm Trekker U tiles (TU004), melted onto the tile and transferred to a pre-chilled 12-well plate, tissue facing upward. Each section was covered with 30 µL of UV cleavage buffer, irradiated for 1 min under a UV lamp (1.2A, cw) and incubated on ice for 7.5 min. Nuclei were released with 200 µL of Buffer A applied four times with gentle repetitive pipetting until dissociation was complete, triturated, washed twice in Buffer B, filtered through a pre-cooled 40 µm strainer, resuspended in 50 µL and counted on a haemocytometer with trypan blue.

Nuclei were processed with the Chromium Next GEM Single Cell 3’ Kit v3.1 and v4 with cell surface protein (10x Genomics) per the manufacturer’s protocol, targeting 20,000 nuclei per sample. Each GEM reaction yielded a whole-transcriptome cDNA library and a spatial barcode library from the photocleaved tags, indexed separately, pooled and sequenced on an Illumina NovaSeq; the shared 10x cell barcode links each nucleus’s coordinates to its transcriptomic profile.

### MERFISH panel design, tissue preparation, and imaging

We designed a custom 500-gene panel (Supplementary Data 1) combining genes implicated in neurogenesis, gliogenesis, neuropeptide signaling and endocrine function with established markers of major brain cell types. Each gene carried a 20-bit binary barcode for decoding sequential imaging data; target sequences came from the Ensembl mouse reference (GRCm38/mm10).

Frozen blocks were coronally sectioned at 10 µm from the preoptic area (approximate Bregma +0.2 mm) onto MERSCOPE slides (Vizgen, 10500001), fixed (4% PFA, 15 min), washed 3× in 1X PBS and permeabilized overnight in 70% ethanol at 4 °C. Using a Vizgen sample preparation kit (10400012), sections were equilibrated in Formamide Wash Buffer (PN 20300002; 30 min, 37 °C), hybridized with the probe set (42-46 h, 37 °C), washed 2× in Formamide Wash Buffer (30 min, 47 °C) and once in Sample Prep Wash Buffer (PN 20300001; 2 min), embedded in a thin polyacrylamide gel (Gel Embedding Premix, PN 20300004) and cleared for 2 to 7 days in 1:100 proteinase K in Clearing Premix (PN 20300003) at 37 °C.

Cleared sections were stained with DAPI and PolyT Staining Reagent (PN 20300021), loaded into the MERSCOPE flow chamber (10000001) and imaged as seven-plane z-stacks at 1.5 µm steps per field of view. The Vizgen MERlin pipeline registered, deconvolved and decoded transcripts, segmented cells with the inbuilt CellPose ’cyto2’ model on DAPI and PolyT signals, and assigned transcripts to cells.

### Xenium tissue and library preparation

Fresh samples were embedded in OCT, frozen, sectioned at 10 µm onto Xenium Slides (10X Genomics) and stored at −80 °C. Sections were fixed and permeabilized per the manufacturer’s protocol and assembled into a Xenium Cassette v2. Priming oligos for the pre-designed Xenium Prime 5K Mouse Pan Tissue & Pathways panel (Supplementary Data 1) were denatured (95 °C, 2 min), diluted in Priming Hyb Buffer to 150 µL per slide, hybridized under a Cassette Insert (50 °C, 90 min; Xenium Thermocycler Adaptor v2), washed in Post-Priming Wash Buffer (PN-2001229; 30 min, 50 °C) and 2× in PBS-Tween (0.05%, PBS-T). After 3× PBS-T, RNase Enzyme in 2X RNase Buffer (20 min, 37 °C) cleaved RNA at the priming site; after 3× 0.5X SSC-T, Polishing Enzyme/Buffer mix (60 min, 37 °C) prepared the template.

Probes were denatured (95 °C, 2 min), diluted in Probe Hyb Buffer B to 150 µL and, after 3× PBS-T, hybridized overnight at 50 °C; unbound probe was removed with 2× PBS-T and Post Hybridization Wash Buffer (15 min, 35 °C). After 3× PBS-T, probe pairs were ligated (30 min, 42 °C) to circularize the probe on the target RNA, then amplified with Amplification Enhancer (2 h, 4 °C; after 3× PBS-T and a 1-min Amplification Enhancer Wash) and Amplification Master Mix (90 min, 30 °C). After 3× TE buffer (pH 8.0), autofluorescence was reduced with Reducing Agent B then Autofluorescence Solution, interspersed with ethanol washes and PBS rehydration. Sections were stored overnight (16–24 h, 4 °C, dark), counterstained with DAPI-containing Xenium Nuclei Staining Buffer, washed 3× in PBS-T and loaded onto the Xenium Analyzer for cyclical probe hybridization, imaging and decoding.

### Data processing and quality control

For Slide-tags, reads were aligned to mm10 with Cell Ranger (v7.1.0, 10x Genomics), retaining intronic reads, and spatial barcodes matched to the bead-array whitelist with CurioTrekker (Curio Bioscience), each nucleus taking the coordinates of the barcodes linked to its 10x cell barcode. Following the original protocol^24^, density-based spatial clustering (DBSCAN) separated ’signal’ barcodes, forming a dense single-nucleus cluster, from dispersed ’noise’; each nuclear coordinate was the UMI-weighted centroid of its signal cluster, and nuclei lacking a clear cluster or with multiple ambiguous clusters were excluded. A second DBSCAN retained only the primary dense cluster, removing nuclei outside the tissue section. In Scanpy we excluded nuclei with a scDblFinder score >0.2, <500 detected genes, <750 UMI counts or >10% mitochondrial reads (Supplementary Fig. 2).

MERlin-generated MERFISH matrices were quality-controlled in Scanpy by removing putative doublets (scDblFinder score >0.2) and cells with <10 detected genes, a volume <100 µm³ or a volume exceeding three times the sample-specific median (∼2000 µm³). For Xenium we removed doublets (score >0.2) and cells with <100 detected genes, <100 total transcripts, an area <10 µm² or an area exceeding three times the sample-specific median (∼250 µm²); the separately captured hemispheres of one control sample were computationally stitched into a single coronal section. For all datasets, counts were normalized to the median of total counts per cell and log-transformed (Supplementary Fig. 2).

### Cell type annotation

To consistently annotate cell types across our MERFISH, Slide-tags, and Xenium datasets, we mapped our data to the Allen Institute’s Mouse Whole Brain Atlas^27^, a single resource that pairs two references under one cell type taxonomy: a spatial reference (MERFISH, 550 genes) registered to the Allen Mouse Brain Common Coordinate Framework v3 (CCFv3), and a matched single-cell RNA-sequencing (scRNA-seq) reference, for which we used only the 10x Genomics Chromium v3 data (2,349,544 cells). From the spatial reference, we selected the four coronal sections (C57BL6J-638850.46–.49) matching our experimental plane, using the anterior commissure and dorsal hippocampus as landmarks. The reference donor was male, but this is unlikely to affect our mapping: the original study identified only a single female-specific cluster, in a subclass absent from our analyses^27^.

To place our cells in a common anatomical coordinate frame, we registered our experimental samples (the query) to the spatial reference using CAST, a graph neural network–based method for spatial transcriptomics^28^. Working with one platform at a time, we restricted the query and the spatial reference to their shared genes and corrected platform-specific expression differences with Scanorama^61^. CAST then learned a shared embedding for each cell from its gene expression and a Delaunay graph of cell positions, using self-supervised training. Guided by these embeddings, each query sample was aligned to a single representative section of the spatial reference (C57BL6J-638850.46) by an affine (global) transformation followed by a non-rigid (free-form deformation) refinement, placing all cells in a shared coordinate system.

We then transferred class and subclass labels to our query cells, combining the spatial reference (anatomical position, but few genes) with the scRNA-seq reference (whole-transcriptome, but no spatial information). We first built a bridge between the references, integrating them into a shared expression space with Scanorama followed by Harmony^61,62^ and linking each spatial reference cell to its 20 nearest scRNA-seq reference cells. We then separately integrated each query dataset with the scRNA-seq reference (Scanorama for platform differences, Harmony for batch effects across samples), so that query and reference cells were comparable by expression (Supplementary Fig. 3A).

To label a query cell, we gathered the spatial reference cells within an adaptive radius of its registered spatial position, followed the bridge to their linked scRNA-seq reference cells, and assigned the label by an inverse-distance-weighted vote over the five closest reference cells by expression. The radius scaled with each dataset’s mean nearest-neighbor distance (100× for the dense MERFISH and Xenium data, 30× for the sparser Slide-tags nuclei), restricting candidates to anatomically plausible identities. Each cell received a class and subclass with three quality metrics: the confidence (the chosen label’s share of the vote), the margin (its lead over the runner-up), and the transcriptomic distance (the cosine distance to the closest matching scRNA-seq reference cell). These spatially-restricted assignments agreed closely with a dissociated, expression-only mapping (94.9% class agreement, 87.9% subclass agreement), with discordance limited to a few anatomically-defined subclasses resolved by the spatial constraint (Supplementary Fig. 3B,C).

We retained only confidently annotated cells. For each dataset, we kept cells with a subclass-assignment confidence ≥ 0.6 and margin ≥ 0.2, and removed spatial and transcriptomic outliers exceeding the per-sample 95th percentile of either the nearest transcriptomic (cosine) distance or the mean distance to their spatial reference neighbors; sample–subclass groups with fewer than 10 cells were also dropped (Supplementary Fig. 2). This yielded 776,646 MERFISH, 70,325 Slide-tags, and 715,429 Xenium cells (Supplementary Data 1). Across platforms, we robustly recovered 84 subclasses spanning 16 classes (each present at ≥100 cells in at least two of the three platforms).

### Correspondence with the published preoptic taxonomy

To relate our subclasses to the preoptic cell types of Moffitt et al. (2018), we labeled their dissociated data (GSE113576) against the same reference^12^. We kept cells carrying their published cluster assignments, excluding those they mark ambiguous or unstable, leaving 30,370 cells in 66 neuronal and 21 non-neuronal clusters. The reference was the scRNA-seq atlas restricted to the anatomical divisions a preoptic dissection can contain (hypothalamus, pallidum, striatum, striatum-like amygdala, olfactory areas and cortical subplate) and capped at 2,000 cells per subclass; we did not restrict it to the subclasses we detect, so that a cluster could map to one absent from our data. Dissociated cells have no coordinates, so labels were transferred by expression alone, with neuronal and non-neuronal cells integrated separately because the two datasets differ in composition. For each compartment we took the shared highly variable genes, corrected them with Scanorama and Harmony as above and computed 30 principal components; each query cell took the majority subclass among its 20 nearest reference cells by cosine similarity, subject to the confidence (≥ 0.6) and margin (≥ 0.2) criteria used for our own annotation, which left 31% of neurons and 1% of non-neuronal cells unassigned. Clusters were scored by the distribution of their assigned cells and counted as detected where at least half fell in the 84 subclasses we report: 44 of 66 neuronal clusters, accounting for 64% of their assigned neurons, and 20 of 21 non-neuronal clusters (Supplementary Fig. 6D).

### Spatial noise correction with ResolVi

Imaging- and bead-based spatial assays misassign transcripts through imperfect segmentation, spillover between neighboring cells and unspecific background, creating spurious co-expression. We corrected each dataset with ResolVi, a deep generative model that decomposes a cell’s observed counts into cell-intrinsic expression, diffusion from neighbors and unspecific background, inferred from the cell’s expression, its neighbors’ expression and the distances between them^31^. We trained one model per dataset on raw counts, treating each sample as a batch and using the 20 nearest spatial neighbors of every cell, in semi-supervised mode with the projected subclass labels so that cell identity informed the decomposition (‘n_latent’ = 10, ‘mixture_k’ = 100; 100 training epochs). From the posterior (30 samples) we estimated the fraction of counts attributable to the true component for each cell and gene, scaled the observed counts by it and rounded to integers. These corrected counts replaced the observed counts downstream.

### Anteroposterior position of sampled sections

To confirm that sections were matched in anteroposterior position across conditions, we estimated the position of every sample against the reference. For each of the 53 reference coronal sections we built a fingerprint from the spatial distribution of its cells: a two-dimensional histogram of cell positions on a fixed 50 × 50 dorsoventral by mediolateral grid, computed separately for each of the 34 cell classes and concatenated. A section’s anteroposterior position was the median x coordinate of its cells in the common coordinate framework. We predicted the position of a query section as the similarity-weighted mean position of its five most similar reference sections, scoring similarity as the cosine between fingerprints.

Leave-one-section-out cross-validation across all 53 reference sections gave a mean absolute error of 0.173 mm overall and 0.074 mm within the window we sampled (4.43 to 6.18 mm, eight sections). We applied the predictor to all 24 profiled sections, using affine-registered coordinates and transferred class labels. Sections were averaged within each of the 21 animals. Predicted positions differed between conditions by about 0.1 mm on each platform (0.08 mm on MERFISH, 0.12 mm on Slide-tags, 0.12 mm on Xenium; Supplementary Fig. 4).

### Differential expression and pathway meta-analysis

We tested for differential expression between reproductive states within each cell type, separately for each platform. ResolVi-corrected counts were aggregated into pseudobulk profiles by sample and cell type, retaining cell types with at least two samples per condition (each pooled from ≥20 cells) and genes detected in ≥30% of profiles within each condition; for Slide-tags, which profiles the whole transcriptome, we kept only protein-coding genes, excluding mitochondrial and ribosomal genes. Each cell type was fit with voomByGroup, a limma-voom variant estimating a separate mean-variance trend per condition, with the log2 number of pooled cells and, for Slide-tags and MERFISH, the log2 pseudobulk library size as covariates. We contrasted pregnant versus nulliparous, postpartum versus pregnant and postpartum versus nulliparous, restricting each test to genes expressed in ≥10 cells in both conditions; postpartum contrasts were run in Slide-tags and MERFISH only, as Xenium lacked a postpartum condition.

To identify changes reproducible across platforms, we combined the per-platform results within each cell type by sum-of-ranks meta-analysis (SumRank)^32^. Genes were ranked within each platform by a signed significance score (the −log10 *p* value times the sign of the log fold change), normalized to [0, 1] with the most upregulated gene at 0, and summed across the platforms whose panels included the gene, requiring at least two. Under the null these ranks are uniform and their sum follows the Irwin-Hall distribution, so a small sum indicates consistent upregulation and a large sum consistent downregulation; we tested each tail separately. Because this analytic null ignores correlation between genes, we calibrated empirically against 1,000 permutations of the condition labels, repeating the per-platform tests and the meta-analysis for each and pooling the statistics within each cell type. A gene’s empirical *p* value per direction was the fraction of this null at least as extreme as observed, and we called genes differentially expressed at empirical *p*≤ 0.05.

Two features of the design bound this calibration. With two to three replicates per condition, the labels admit 20 distinct assignments for Slide-tags and MERFISH and 6 for Xenium, so the smallest attainable per-platform permutation *p* value is 0.10, 0.10 and 0.33; the 1,000 permutations sample these repeatedly estimating the null precisely rather than extending its range. We therefore treat empirical *p* ≤ 0.05 as an exploratory threshold nominating candidates, and rest our conclusions on reproducibility (a change is reported only where at least two independently acquired platforms measured the gene and ranked it consistently, D ≥ 2), coordination (we interpret sets of genes and pathways moving together across related cell types rather than individual gene calls) and orthogonal protein-level validation (Figs. 3E, 5E). Benjamini-Hochberg correction of these empirical *p* values leaves no gene and cell type change below a false discovery rate of 0.10 in pregnancy; the smallest is 0.19 (Supplementary Data 3).

The same procedure was carried through to pathway enrichment. Genes were ranked per platform and cell type by the signed score and tested with fgsea (minimum gene-set size 15) against 3,043 Gene Ontology biological-process gene sets spanning 14 themes relevant to maternal-brain physiology, among them reproduction, hormonal signaling, glia and myelination, synaptic plasticity, vasculature and metabolism (Supplementary Data 4). Pathways were ranked by a signed enrichment score (the −log10 *p* value times the sign of the normalized enrichment score), summed as for genes and calibrated against fgsea run on each permutation’s results. Pathways were considered enriched at empirical *p* ≤ 0.05. As for genes, Benjamini-Hochberg correction of these empirical *p* values leaves no pathway and cell type pair below a false discovery rate of 0.10 (smallest 0.15; Supplementary Data 4). This is therefore also an exploratory threshold, and we interpret pathways where they recur across related cell types and reproduce across platforms.

### Intercellular signaling analysis

We characterized spatially resolved cell-cell communication in the Slide-tags and Xenium datasets with LIANA^63^, using a ligand-receptor resource combining LIANA’s mouse-consensus database with the neurotransmitter and neuropeptide interactions in NeuronChat ^64^. For each sample we built a spatial neighbor graph with a Gaussian kernel (bandwidth 100 µm for Slide-tags, 50 µm for Xenium), setting each cell’s signaling neighborhood, and kept only spatially structured signals with a Moran’s I filter in squidpy at two stages^65^: first on log-normalized expression (I > 0.01, FDR < 0.05), from which we computed a per-cell inflow score for each ligand-receptor pair — the cell’s receptor expression weighted by the distance-weighted ligand expression of its neighbors — and again on the resulting interaction scores (I > 0.01, FDR ≤ 0.05). Retained scores were aggregated to source and target cell type pairs and each interaction’s specificity tested against a null of 1,000 cell type-label permutations, recording its mean score and specificity *p* value. Per-sample results were averaged within condition and signaling differences taken as the change in this mean score (all three contrasts for Slide-tags; pregnant versus nulliparous for Xenium). For pregnant versus nulliparous we focused on a curated set of ligand-receptor pairs grouped by signaling theme and required reproducibility: we kept interactions shifting in the same direction on both platforms, each exceeding that platform’s median effect size, and took the mean of the two changes as the combined effect. Interactions measurable on only one platform, for example pairs outside the Xenium gene panel, were retained from Slide-tags alone at the top decile of effect magnitude (Supplementary Data 5).

### Differential abundance and proximity analyses

We asked whether cell type composition shifted across reproductive states. For each dataset we tabulated subclass counts per sample and modeled their relative abundance with the crumblr R package^66^, which applies a centered log-ratio transformation with observation-level precision weights, fit with the dream function from variancePartition with condition as a fixed effect. We tested the same three contrasts as for differential expression at a Benjamini-Hochberg false discovery rate (FDR) ≤ 0.10. No subclass reached an FDR of 0.10 on any platform, and the few nominal shifts did not agree in direction across platforms (Supplementary Fig. 5; Supplementary Data 2).

We separately tested local co-localization in the imaging-based MERFISH and Xenium datasets, whose single-cell coordinates support neighborhood statistics. For each cell we counted neighbors of every subclass within 20 times the sample’s median nearest-neighbor distance (∼200 µm) and converted the counts to enrichment z-scores against a null from 100 random permutations of cell type labels. Per cell type pair, z-scores were averaged per sample and compared between conditions with a weighted linear model (limma), weighting by contributing cells and restricting to pairs co-occurring in at least five cells per sample (FDR ≤ 0.10 per dataset). MERFISH and Xenium results for pregnant versus nulliparous were combined by the same meta-analysis and permutation calibration as for differential expression (Supplementary Data 2).

### Genetically informed spatial mapping of cells for complex traits (gsMap)

We collected GWAS summary statistics for 16 human neurological and psychiatric traits and disorders from the GWAS Catalog (Supplementary Data 8) and applied gsMap^46^ to the Slide-tags and Xenium datasets, separately for each reproductive state, mapping mouse genes to their human homologs. gsMap assigns each gene a local, spatially aware Gene Specificity Score in each cell with a graph neural network, maps these scores to nearby SNPs and applies stratified LD score regression (S-LDSC) to test each cell for heritability enrichment, yielding a per-cell association P value per trait. We combined these within each cell type by Cauchy combination, giving a per-condition enrichment for each cell type-trait pair (Benjamini-Hochberg FDR across trait × cell type pairs).

To test whether this enrichment shifts across states, we summarized each dataset, trait and cell type as the per-sample mean of the cell-level association scores (−log10 P), retaining sample groups with at least 20 cells, and tested for a difference with a Welch t-test where both conditions had at least two such samples. Slide-tags and Xenium were combined for pregnant versus nulliparous by the same meta-analysis and label-permutation calibration as for differential expression; postpartum contrasts were tested in Slide-tags alone (Supplementary Data 8).

### Immunohistochemistry validation experiments

Mice were anesthetized with isoflurane and intracardially perfused with PBS then 4% PFA in phosphate buffer. Brains were post-fixed in 4% PFA overnight at 4 °C, dehydrated in 30% sucrose for >48 h at 4 °C, embedded in OCT compound (Fisher Healthcare, #4585) and cryosectioned at 15 µm onto Superfrost Plus slides (Fisher Scientific, #1255015).

For FKBP5 staining only, antigen was retrieved in Tris-EDTA buffer (10 mM Tris base, 1 mM EDTA, 0.05% Tween 20, pH 9.0) at 95–100 °C on a heat block for 20 min, followed by 10 min cooling under running cold water. Sections were blocked and permeabilized (3% bovine serum albumin, 0.3% Triton X-100 in 1x PBS; 1 h, room temperature), incubated with primary antibodies in blocking buffer overnight at 4 °C, washed 3× in PBST (0.1% Tween 20 in 1x PBS), incubated with secondary antibodies and 1 µg/mL DAPI (Thermo Fisher, #62248) in PBST (2 h, room temperature), washed 3× in PBST and mounted in ProLong Gold Antifade Mountant (Fisher Scientific, #P36930). Primary antibodies were goat anti-ApoE (Sigma-Aldrich, #AB947, 1:200), mouse anti-IBA1 (Abcam, #ab283319, 1:100), rabbit anti-NG2 (Sigma-Aldrich, #AB5320, 1:200), chicken anti-S100β (Aves Labs, #S100B-0100, 1:500), rabbit anti-HMGCR (Proteintech, #13533-1-AP, 1:100), rabbit anti-FKBP5 (Proteintech, #14155-1-AP, 1:50) and goat anti-Mertk (R&D Systems, #AF591, 1:100). Secondary antibodies, all 1:500 from Thermo Fisher, were donkey anti-goat IgG (H+L) Alexa Fluor 594 (#A-11058), donkey anti-mouse IgG (H+L) Alexa Fluor 594 (#A-21203), donkey anti-mouse IgG (H+L) Alexa Fluor 488 (#A-21202), donkey anti-chicken IgY (H+L) Alexa Fluor 488 (#A-78948) and donkey anti-rabbit IgG (H+L) Alexa Fluor 647 (#A-31573). Sections were imaged on an LSM880 confocal microscope with ZEN Black (2.3 SP1, Zeiss) and analyzed in the FIJI version of ImageJ (1.54p with Java 1.8.0_452, 64 bit; NIH).

We validated selected protein changes in a separate cohort of nulliparous and pregnant mice (n = 7 per group), quantifying APOE and HMGCR within defined cell types and FKBP5 across the striatum. APOE and HMGCR signal counted as cell type-specific only where it overlapped the corresponding marker: IBA1 for microglia and NG2 for oligodendrocyte precursor cells for APOE, S100β for astrocytes for HMGCR. To confirm that the target lay within the marked cell rather than adjacent to it, we acquired z-stacks of five slices at 1 µm intervals and assessed localization in three dimensions. Colocalization was quantified with the DiAna (Distance Analysis) plugin in Fiji^67^: target and marker objects were segmented independently by DiAna object-based segmentation, overlapping marker objects identified with its colocalization function, and the integrated density of target protein within marker-positive objects normalized to total marker-positive area. FKBP5 was assessed regionally instead: we imaged the striatum, applied a single intensity threshold across all images and measured FKBP5-positive area within a striatal region of interest in Fiji. DAPI was the nuclear counterstain throughout. We analyzed three to five images per mouse, used the per-mouse mean as the biological replicate and compared groups with an unpaired two-sided Welch t-test (Supplementary Data 6).

### Vascular microscopy validation experiments

Wild-type C57BL/6 mice (7 weeks old; The Jackson Laboratory) were studied in two female cohorts: nulliparous, group-housed to synchronize estrous cycles, and timed-pregnant, harvested at embryonic day 17.5 to 18.5 (four and five animals; one further nulliparous animal was excluded for unusable sections). Mice were transcardially perfused with 10 mL of phosphate-buffered saline (PBS) then 10 mL of 4% paraformaldehyde, each over 10 min; brains were cryoprotected through a 10%, 20% and 30% sucrose gradient, embedded in 1:1 30% sucrose and OCT, flash-frozen and cut at 30 µm onto glass slides.

Sections were rehydrated in PBS, blocked and permeabilized (10% fetal bovine serum, 3% bovine serum albumin, 1% glycine, 0.4% Triton X-100 in TBS; 1 h, room temperature), incubated overnight at 4 °C with rat anti-mouse CD31 (1:200; BD Pharmingen, 557355), washed 3× for 10 min in PBST, incubated for 90 min with Alexa Fluor 555-conjugated goat anti-rat IgG (1:500; Invitrogen, A-21434) and mounted in DAPI Fluoromount-G (SouthernBiotech, 0100-20). We imaged the preoptic area on a Leica Stellaris 5 confocal microscope (40x oil-immersion, 30 µm z-stacks at 0.7 µm steps), optimizing detector gain per sample, so intensity is not comparable between images and only vessel geometry was analyzed. In FIJI we made maximum-intensity projections of the CD31 and DAPI channels, subtracted background with an 80-pixel rolling ball, despeckled and thresholded CD31 manually to a binary vessel mask, cropped so that fields from one animal do not overlap: 12 nulliparous and 16 pregnant fields at 1.7583 pixels per µm.

Morphometrics were computed in Python (scikit-image, skan, SciPy). Fields contain off-section space and being larger in nulliparous mice, cannot serve as a denominator; we recovered the analyzed region per image as the morphological closing of the vessel mask at 100 µm and counted inside a further 25 µm erosion, computing skeletons and distance transforms on the full mask so the boundary neither truncates branches nor inflates endpoint counts. Objects and holes below 20 µm² were removed. From each mask we measured vessel area fraction, skeleton length density, junction, branch and endpoint density, mean branch length and tortuosity, lacunarity and fractal dimension, vessel diameter from the distance transform, and tissue distance to the nearest vessel; from DAPI, nuclear density and the density of nuclei within 3 µm of a vessel, which in 30 µm projections is relative rather than absolute. Re-segmenting every image by Li thresholding controlled for the manual threshold (mean Dice coefficient 0.93 across 28 fields).

Fields are technical replicates, so we averaged them within animal and tested at the animal level, where the variance lies (vessel-area intraclass correlation 0.85). Each of the eighteen measures was compared with a Welch t-test, an exact permutation test over all 126 assignments of nine animals to groups of four and five, and a linear mixed model on all 28 fields with a random intercept per animal, corrected by FDR. Because a null result at this sample size is a statement about precision, we report the confidence interval of every difference and the minimum detectable effect at 80% power. Results were unchanged under alternative denominators, despeckling thresholds, domain radii and thresholding rules, and on removal of any single animal (Supplementary Fig. 7; Supplementary Data 7).

## Data availability

Raw sequencing data (FASTQ) and processed files (.h5ad) for the Slide-tags dataset have been deposited in the Gene Expression Omnibus (GEO) under accession code GSE313279. Processed data (.h5ad) for the MERFISH dataset has been deposited in GEO under accession code GSE314151. Processed data for the Xenium dataset (.h5ad) has been deposited in GEO under the accession code GSE342797.

## Code availability

Custom code used for the analyses in this study are available on GitHub at https://github.com/keon-arbabi/spatial-pregnancy

## Acknowledgements

This work was supported by the Centre for Addiction and Mental Health (CAMH) Discovery Fund and the Krembil Foundation (to S.J.T.). S.J.T. acknowledges support from the Natural Sciences and Engineering Research Council of Canada (NSERC) (RGPIN-2020-05834 and DGECR-2020-00048), the Canadian Institutes of Health Research (CIHR) (PJT-191747, NGN-171423, and PJT-175254), and a Brain Canada Future Leaders in Canadian Brain Research grant. B.T.K. acknowledges support from the Hospital for Sick Children (SickKids), the SickKids Foundation, as well as the Merton Bernfield Chair in Neonatology and the Division of Newborn Medicine at Boston Children’s Hospital. The facility is supported by the Canada Foundation for Innovation and the Ontario Government. L.A.M.G. is supported by a womenmind grant (CAMHF-1197), the New Frontiers in Research Fund (NFRF-T-2022-00051), a Brain Canada Basics of Better Mental Health Program grant and the Women’s Health Research Cluster. M.Y.F. is supported by a CIHR Canada Graduate Scholarship and an Ontario Graduate Scholarship. K.A. is supported by a Koerner Graduate Award.

## Author contributions

B.T.K. conceptualized the study. B.K. carried out MERFISH, Slide-tags, and Xenium sample preparation and data generation. K.A., S.G., and H.H. performed the formal analysis. K.A. created the visualizations. K.A. wrote the original draft. K.A., S.G., M.E.F., B.K., L.A.M.G., S.J.T., and B.T.K. reviewed and edited the manuscript. S.J.T. and B.T.K. supervised the work.

## Description of Additional Supplementary Files

**File Name:** Supplementary Data 1

**Description:** Sample metadata and post-quality-control cell counts for the three platforms, the MERFISH and Xenium gene panels annotated with the cell subclasses each gene discriminates and its pathway themes, and source data for the marker dot plot. Related to Figure 1.

**File Name:** Supplementary Data 2

**Description:** Differential cell type abundance between reproductive states, per platform and compared across platforms (centered log-ratio transformation with precision weights, two-sided moderated t-test, Benjamini-Hochberg correction), and differential local co-occurrence of cell subclass pairs on MERFISH and Xenium, per platform and combined by sum-of-ranks meta-analysis (two-sided moderated t-test on permutation-derived enrichment z-scores; one-sided empirical P values per direction, Benjamini-Hochberg correction). Includes per-sample source data and a sheet defining every column. Related to Supplementary Figure 5.

**File Name:** Supplementary Data 3

**Description:** Differential expression between reproductive states per platform, cell subclass and gene (pseudobulk, voomByGroup and limma, two-sided moderated t-test, Benjamini-Hochberg correction), the cross-platform sum-of-ranks meta-analysis calibrated against 1,000 permutations of the condition labels (one-sided empirical P values per direction), and the genes changed per cell subclass shown in Figure 2A. Restricted to the subclass and gene pairs entering the meta-analysis. Related to Figure 2A.

**File Name:** Supplementary Data 4

**Description:** Gene set enrichment against 3,043 curated GO biological processes per platform, cell subclass and pathway (fgsea on genes preranked by signed significance), the cross-platform sum-of-ranks meta-analysis calibrated against 1,000 permutations of the condition labels (one-sided empirical P values per direction), and the keyword rules assigning pathways to themes. Restricted to the subclass and pathway pairs entering the meta-analysis; the unrestricted table of all tests is deposited with the study data. Related to Figures 2B and 3–5.

**File Name:** Supplementary Data 5

**Description:** Intercellular signaling in pregnancy. Ligand-receptor interactions scored per cell with LIANA on a spatial neighborhood graph and aggregated to ordered pairs of cell subclasses, reported for every interaction changing consistently on both platforms (measured on Slide-tags and Xenium, same direction on each, and at least the median absolute change of that platform), for the curated ligand-receptor pairs of the chord panels, and for the ligand-receptor resource searched. No P value is reported for a difference: the permutation test establishes that an interaction is specific to a pair of cell types, not that it changes between conditions, so agreement across two independently acquired platforms is the criterion. Related to Figures 3F, 4E and 5F.

**File Name:** Supplementary Data 6

**Description:** Immunofluorescence validation of four protein changes in nulliparous and pregnant mice, seven per group: FKBP5 across the striatum, and APOE within microglia (IBA1) and oligodendrocyte precursor cells (NG2) and HMGCR within astrocytes (S100β). Reports every quantified image, the per-mouse means that are the biological replicates, and the group comparisons (unpaired two-sided Welch t-test on per-mouse means). Related to Figures 3E and 5E.

**File Name:** Supplementary Data 7

**Description:** Quantitative morphometry of the CD31-stained vascular bed in the preoptic area, in an independent cohort of four nulliparous and five pregnant mice (12 and 16 imaged fields). Reports 18 measures spanning vessel density, network architecture, caliber, tissue supply and cellularity, each compared between groups on per-animal means (unpaired two-sided Welch t-test, exact permutation test over all 126 assignments, and a linear mixed model over fields with animal as a random effect; Benjamini-Hochberg correction across the 18 measures), together with the per-animal and per-field measurements, the smallest change the cohort could have detected, and every analytic choice varied in turn. Related to Figure 4F and Supplementary Figure 7.

**File Name:** Supplementary Data 8

**Description:** Heritability enrichment of 16 human neurological and psychiatric traits across the maternal brain, computed with gsMap on the Slide-tags and Xenium datasets in each reproductive state. Reports the source genome-wide association study for each trait, the enrichment of each trait in each cell subclass per platform and state (Cauchy combination of per-cell association P values, Benjamini-Hochberg correction), the change in enrichment between reproductive states per platform (Welch t-test on per-sample means), the Slide-tags and Xenium sum-of-ranks meta-analysis of the pregnant versus nulliparous contrast calibrated against label permutations, and the per-sample association scores these rest on. Related to Figure 6.

