## Supplementary Figures for "Spatial mapping of cellular and molecular plasticity in the maternal and postpartum mouse brain"

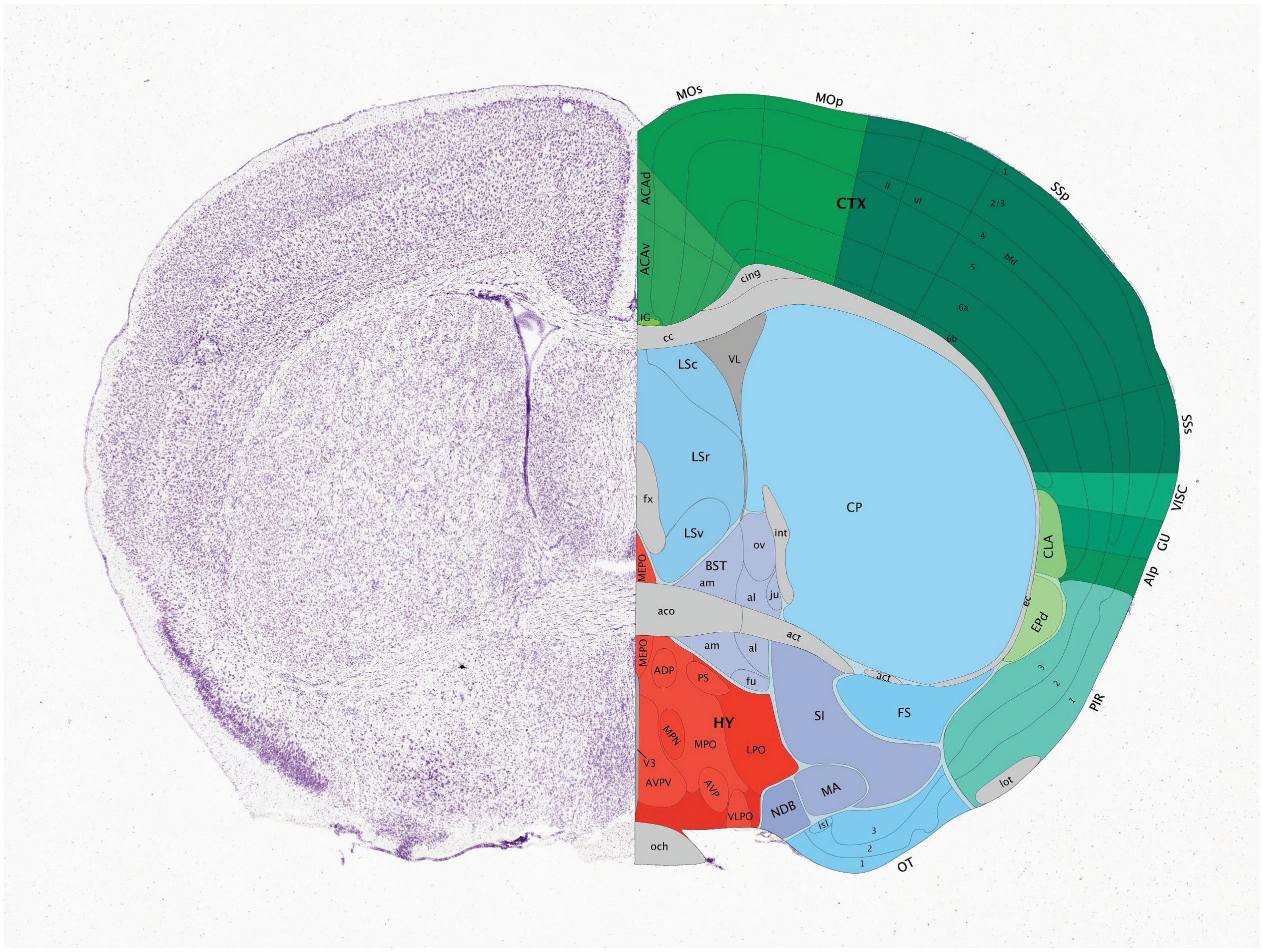


**Supplementary Fig. 1.** ***Anatomical reference for the sampled coronal plane.*** Nissl-stained hemisphere (left) and the corresponding delineated annotation from the Allen Reference Atlas (right) at the coronal level sampled in this study (approximately Bregma +0.2 mm). The four regions profiled appear as the isocortex (CTX), caudoputamen (CP), lateral septal complex (LSc, LSr, LSv) and preoptic hypothalamus (HY). Cortex: ACAd, ACAv, anterior cingulate area, dorsal and ventral parts; MOp, MOs, primary and secondary motor areas; SSp, SSs, primary and supplemental somatosensory areas, with SSp layers (1, 2/3, 4, 5, 6a, 6b) and the lower limb (ll), upper limb (ul) and barrel field (bfd) subfields outlined; GU, gustatory area; VISC, visceral area; AIp, agranular insular area, posterior part; PIR, piriform area (layers 1-3); CLA, claustrum; EPd, endopiriform nucleus, dorsal part; IG, induseum griseum. Striatum and pallidum: CP, caudoputamen; LSc, LSr, LSv, lateral septal nucleus, caudal, rostral and ventral parts; BST, bed nuclei of the stria terminalis (al, anterolateral; am, anteromedial; fu, fusiform; ju, juxtacapsular; ov, oval); FS, fundus of striatum; MA, magnocellular nucleus; NDB, diagonal band nucleus; SI, substantia innominata; OT, olfactory tubercle (layers 1-3; isl, islands of Calleja). Hypothalamus: MPO, medial preoptic area; MPN, medial preoptic nucleus; LPO, lateral preoptic area; MEPO, median preoptic nucleus; ADP, anterodorsal preoptic nucleus; AVP, anteroventral preoptic nucleus; AVPV, anteroventral periventricular nucleus; VLPO, ventrolateral preoptic nucleus; PS, parastrial nucleus. Fiber tracts and ventricles: aco, anterior commissure, olfactory limb; act, anterior commissure, temporal limb; cc, corpus callosum; cing, cingulum bundle; ec, external capsule; fx, fornix; int, internal capsule; lot, lateral olfactory tract; och, optic chiasm;

VL, lateral ventricle; V3, third ventricle. Adapted from the Allen Mouse Brain Atlas (Allen Institute for Brain Science; atlas.brain-map.org).

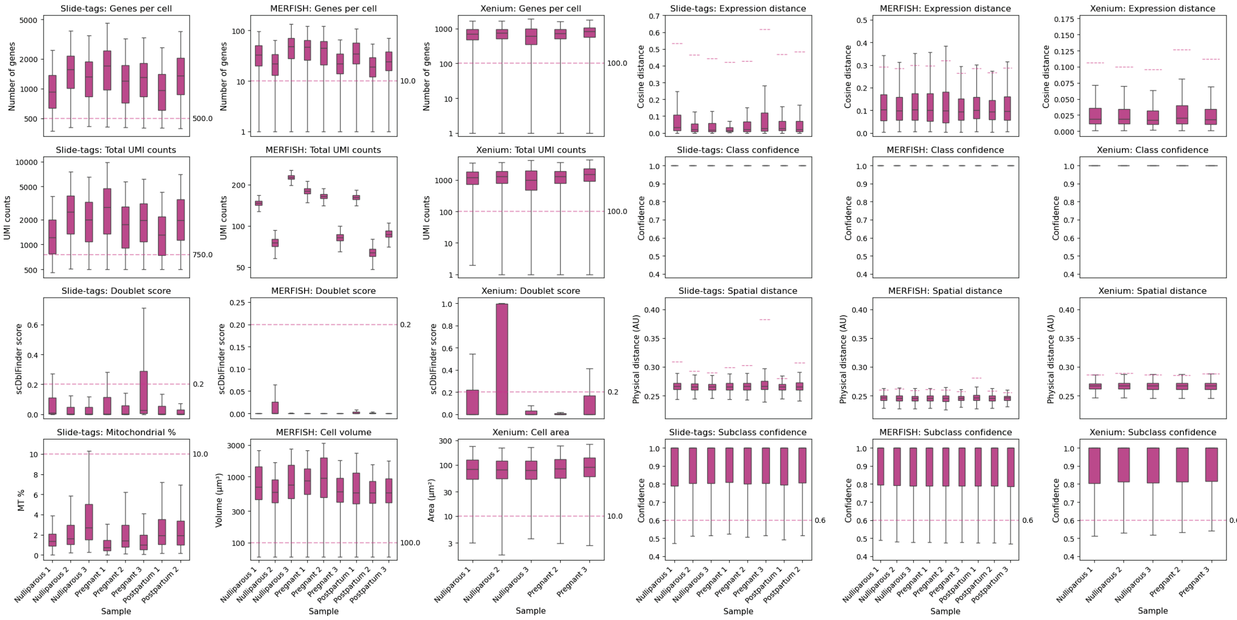


**Supplementary Fig. 2.** ***Quality control and cell type annotation performance for the Slide-tags, MERFISH and Xenium spatial datasets.*** Each box shows the per-cell metric for one biological replicate. The three left-hand columns report pre-filtering quality-control metrics for each platform: detected genes per cell, total UMI counts, per-cell doublet score (scDblFinder) and a cell-size metric — mitochondrial read percentage (MT %) for Slide-tags, cell volume (µm³) for MERFISH and cell area (µm²) for Xenium. Pink dashed lines mark the exclusion thresholds applied during preprocessing: Slide-tags, >500 genes, >750 UMIs, doublet score <0.2 and MT <10%; MERFISH, >10 genes, doublet score <0.2 and cell volume >100 µm³; Xenium, >100 genes, >100 UMIs, doublet score <0.2 and cell area >10 µm². For MERFISH and Xenium, cells were additionally capped at 3× the per-sample median volume/area. The three right-hand columns report cell type annotation quality after CAST integration with the Allen Mouse Brain Atlas: the cosine distance between query and reference expression profiles (Expression distance), the confidence of broad cell-class assignment (Class confidence), the physical distance to the mapped reference location (Spatial distance) and the confidence of fine-grained subclass assignment (Subclass confidence). Pink dashed lines indicate the criteria used to retain high-confidence annotations: a global subclass-confidence cutoff (>0.6; horizontal line) together with per-sample 95th-percentile limits on expression and spatial distance (per-sample segments), above which cells were discarded; an additional subclass-margin criterion (>0.2) was applied but is not shown. Class confidence is displayed for reference and was not used for filtering. Samples comprise Nulliparous, Pregnant and Postpartum replicates; the Xenium cohort includes Nulliparous and Pregnant replicates only.

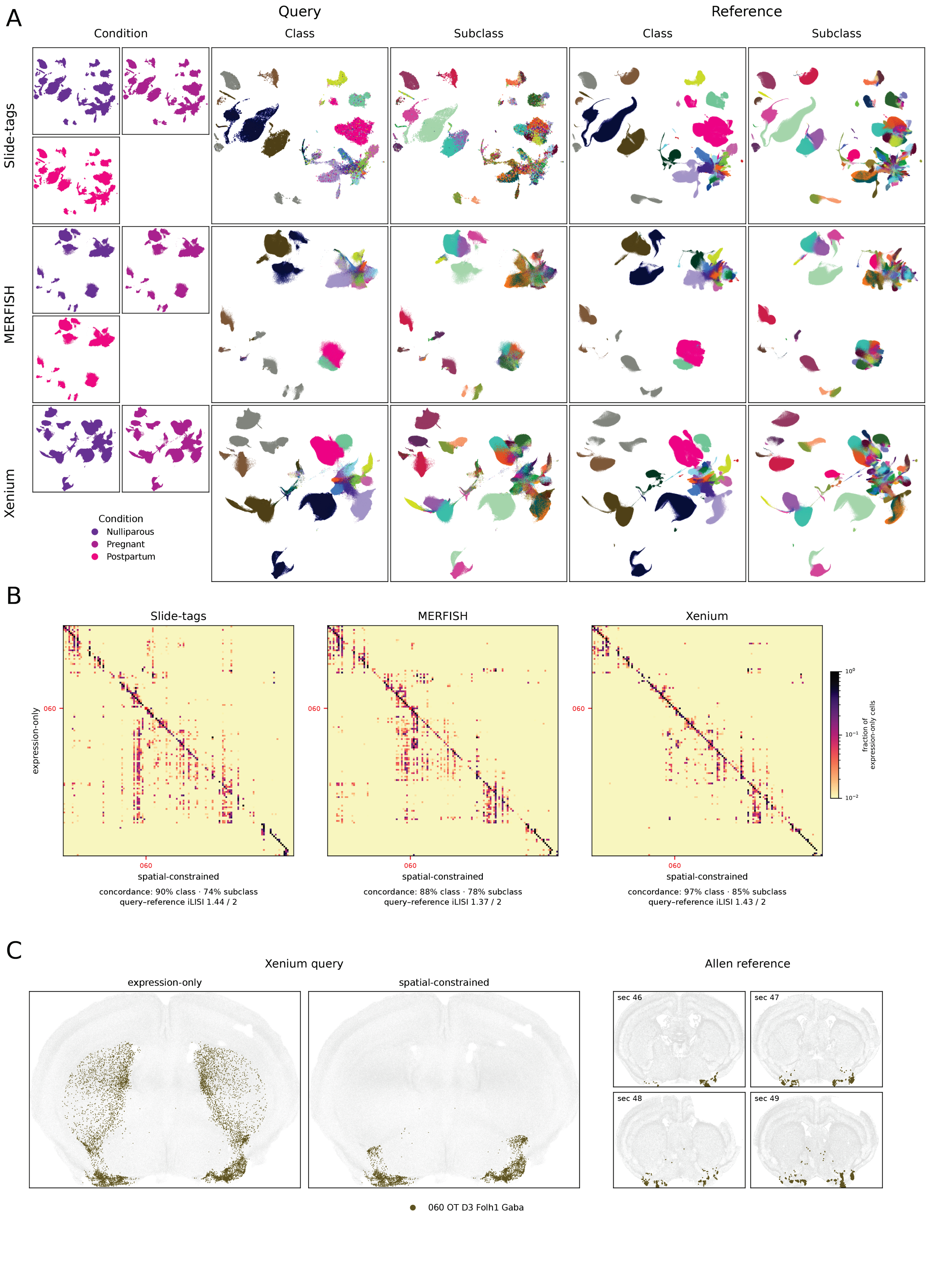
A
Query
Reference
Condition
Class
Subclass
Class
Subclass
Slide-tags
MERFISH
Xenium
Condition
Nulliparous
Pregnant
Postpartum
B
Slide-tags
MERFISH
Xenium
expression-only
060
060
060
060
060
060
spatial-constrained
spatial-constrained
spatial-constrained
concordance: 90% class · 74% subclass
query–reference iLISI 1.44 / 2
concordance: 88% class · 78% subclass
query–reference iLISI 1.37 / 2
concordance: 97% class · 85% subclass
query–reference iLISI 1.43 / 2
fraction of expression-only cells
C
Xenium query
expression-only
spatial-constrained
Allen reference
sec 46
sec 47
sec 48
sec 49
060 OT D3 Folh1 Gaba

**Supplementary Fig. 3.** ***Shared-latent-space projection assigns Allen atlas cell types consistently across platforms and is refined by spatial constraint.*** **A)** For each spatial platform (Slide-tags, MERFISH, Xenium; rows), query cells and the scRNA-seq reference were co-embedded in a shared Harmony latent space and visualized with a single combined UMAP per platform, so query and reference occupy the same coordinates. Query cells (left three columns) are colored by condition, cell class, and subclass; the reference (right two columns) by class and subclass. In the condition column each condition is shown in its own quadrant (nulliparous, purple; pregnant, magenta; postpartum, pink; Xenium lacks a postpartum sample). Class and subclass colors follow the Allen atlas palette. Query and reference cells populate matching regions of every embedding, indicating successful cross-modality integration. **B)** Row-normalized confusion between expression-only (unconstrained) subclass labels (rows) and spatial-constrained labels (columns) for each platform; color is the fraction of a row's expression-only cells falling in each column (log scale), so a dark diagonal denotes agreement. Beneath each matrix: label agreement between the two schemes (percent of cells with matching class and subclass) and query–reference integration (iLISI; 1 = fully separated, 2 = fully mixed). The featured subclass 060 OT D3 Folh1 Gaba is boxed in red and marked on both axes. **C)** Spatial distribution of 060 OT D3 Folh1 Gaba (olive) in a representative Xenium coronal section, comparing expression-only and spatial-constrained assignments, alongside the same subclass in the Allen reference atlas across four coronal sections (sec 46–49). Expression-only labelling over-calls this subclass broadly across the section, whereas the spatial constraint confines it to the olfactory tubercle, matching the reference; all other cells are shown in grey.

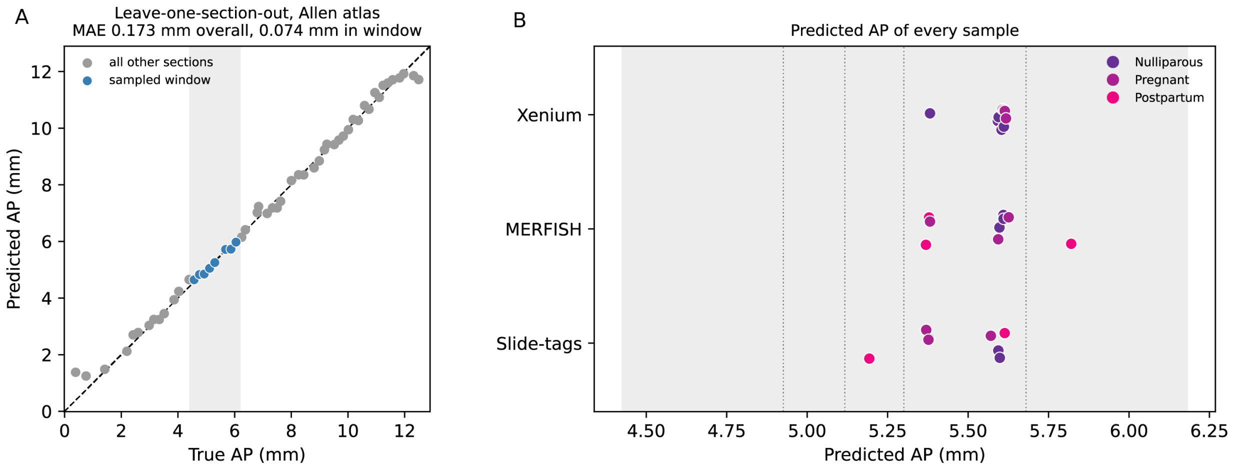


**Supplementary Fig. 4.** ***Sections sampled for each condition were cut at matched anteroposterior positions.*** Position was estimated by matching the spatial distribution of cell classes against the Allen reference: for each of the 53 reference coronal sections a fingerprint was built as a two-dimensional histogram of cell positions on a fixed 50 × 50 dorsoventral by mediolateral grid, computed per cell class and concatenated, and a query section was assigned the similarity-weighted mean position of its five most similar reference sections (cosine similarity between fingerprints; Methods). **A)** Leave-one-section-out cross-validation on the reference. Predicted against true anteroposterior position for every reference section, with the line of identity dashed. The shaded band marks the window sampled in this study (4.43 to 6.18 mm); sections falling within it are blue and all others gray. Mean absolute error was 0.173 mm across all 53 sections and 0.074 mm across the eight within the sampled window. **B)** Estimated position of every animal, by platform (rows) and reproductive state (color), computed from affine-registered coordinates and transferred class labels. Position was estimated separately for each section and averaged within an animal where more than one section was profiled, so each point is one mouse. Dotted lines mark the four reference sections spanning the sampled plane and the shaded band is as in A; points are jittered vertically for visibility. Mean position differed between conditions by about 0.1 mm on every platform (0.08 mm on MERFISH, 0.12 mm on Slide-tags, 0.12 mm on Xenium).

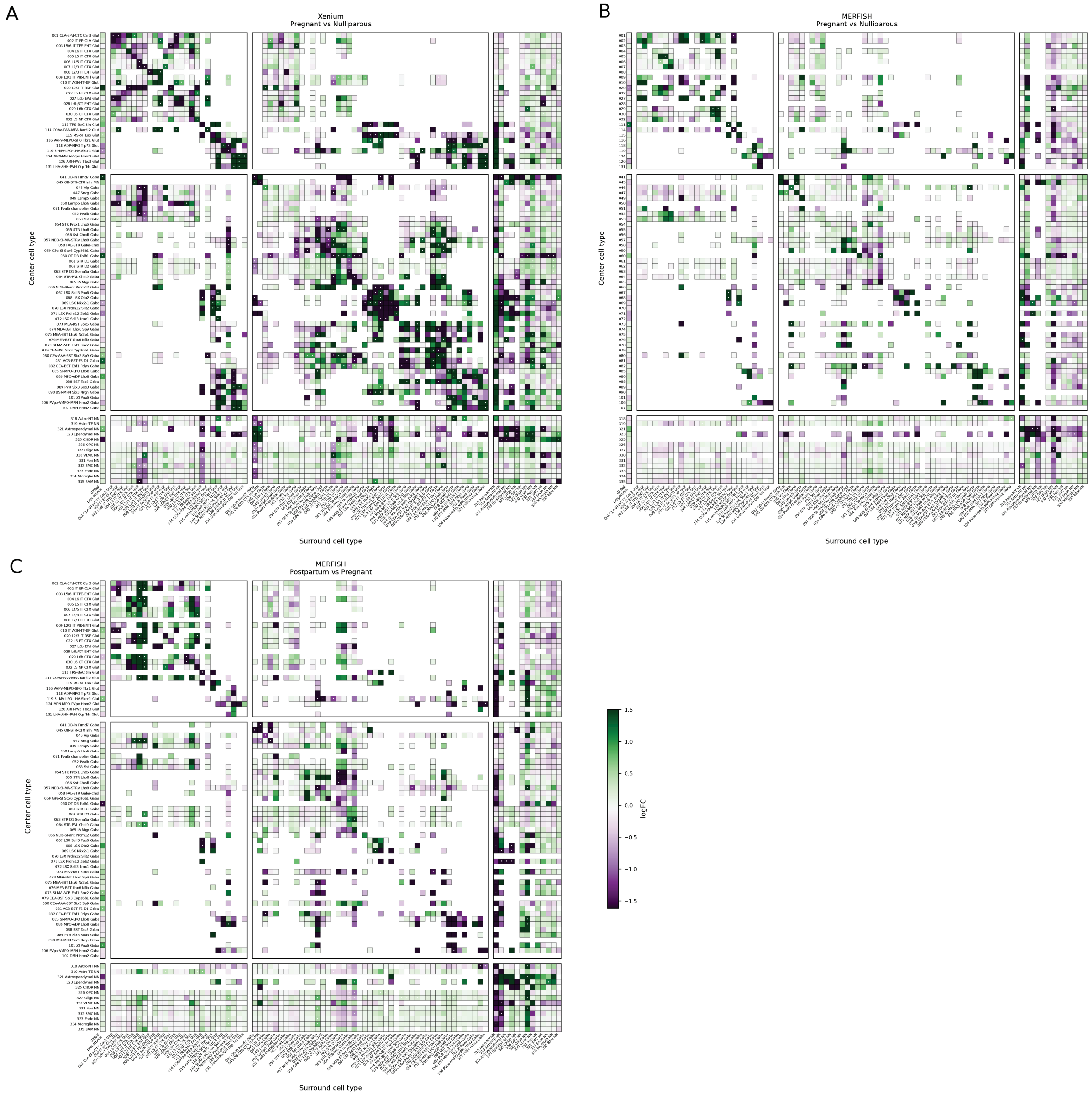


**Supplementary Fig. 5.** ***Cell type spatial co-occurrence and composition shift only sparsely across the peripartum window, without reproducible reorganization.*** For the 84 subclasses robustly detected across platforms, spatial co-occurrence was scored per platform as the log fold-change in how enriched or depleted each surround subclass is within the local neighborhood (~200 μm) of each center subclass, relative to permuted cell type labels; the adjacent single column gives each subclass's change in global proportion (overall abundance). In every heatmap, rows are center subclasses and columns are surround subclasses, both grouped into glutamatergic, GABAergic, and non-neuronal blocks. Fill color is the log fold-change (green, increased; purple, decreased co-occurrence or proportion in the later condition of each contrast; clipped at the 5th–95th percentiles); glyphs mark per-platform significance (*, FDR < 0.10; •, nominal p < 0.05). **A)** Xenium and **B)** MERFISH, pregnant versus nulliparous; **C)** MERFISH, postpartum versus pregnant (postpartum was not profiled by Xenium). No proximity shift survived multiple-testing correction on any platform (no pairs at FDR < 0.10), and the nominally significant pairs were scattered across cell types rather than concentrated in any population – consistent with pregnancy remodeling the maternal brain within its existing cellular architecture rather than through coordinated spatial rearrangement.

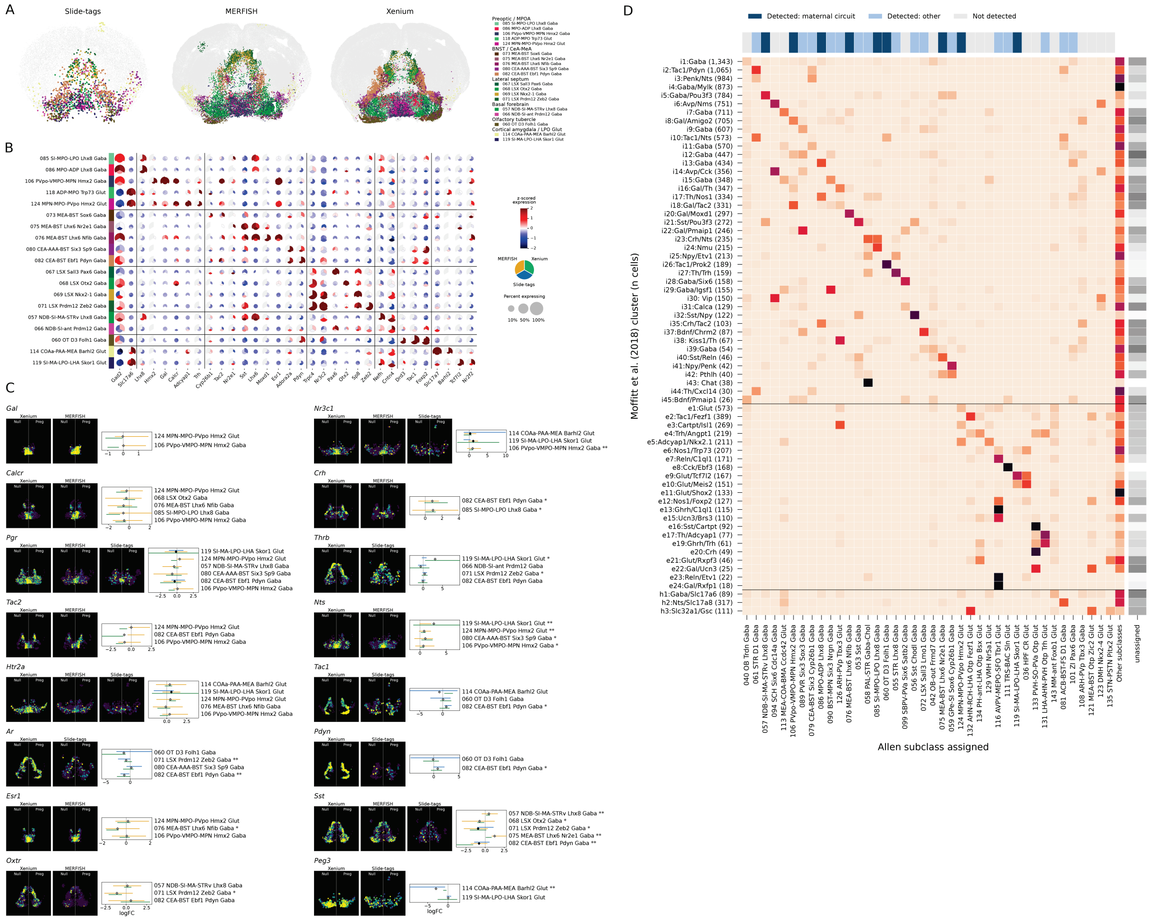

**Supplementary Fig. 6.** ***Maternal-control circuits are captured on all three platforms and keep their identity in pregnancy, apart from a focused neuropeptidergic program.*** Nineteen subclasses constituting the core maternal-behavior circuitry – preoptic hypothalamus (MPOA), extended amygdala (BNST, CeA-MeA), lateral septum, basal forebrain, olfactory tubercle and cortical amygdala – were examined across Slide-tags, MERFISH and Xenium, with differential expression for the pregnant versus nulliparous contrast meta-analyzed as in Fig. 2. **A)** Spatial distribution of the nineteen subclasses. All pregnant sections are superimposed per technology (Slide-tags, left; MERFISH, middle; Xenium, right); cells of these subclasses are colored by subclass, grouped into six anatomical families, and all other cells are gray. **B)** Identity-marker dot plot confirming the annotations: the nineteen subclasses (rows, grouped by family) against 32 canonical markers (columns, grouped by the family they label), with three wedges per dot, one per technology, encoded as in Fig. 1C. **C)** Gene cards contrasting stable maternal effectors with pregnancy-regulated neuroendocrine genes. Effectors that are expressed but transcriptionally stable (*Gal, Calcr, Pgr, Tac2, Htr2a*) are shown alongside the genes that change: the neuropeptides Crh and Nts increase across several subclasses, the androgen receptor Ar decreases and the thyroid-hormone receptor Thrb increases in two each, and the remaining genes (*Esr1, Oxtr, Nr3c1, Tac1, Pdyn, Sst, Peg3*) change sparsely or inconsistently in direction. For each gene, single-cell expression is mapped in nulliparous (left of midline) and pregnant (right) hemispheres for every technology assaying it, beside a forest plot of per-platform and meta-analyzed log2 fold change (95% CI) across the cell types expressing it; asterisks on cell type labels denote meta significance (*, P <= 0.05; **, P <= 0.01; ***, P <= 0.001). **D)** Correspondence between the preoptic cell types of Moffitt et al. (2018) and the subclasses used here. Their dissociated preoptic cells were labeled with Allen subclasses by expression alone, without the spatial constraint used elsewhere and without restricting the reference to the subclasses we detect, so that a cluster could correspond to one absent from our data (see Methods). Rows are their 66 neuronal clusters, grouped into inhibitory, excitatory and hybrid blocks with cell numbers in parentheses; columns are the 40 subclasses taking the largest share of assigned cells, with the remainder pooled as "Other subclasses" so that each row sums to one. Fill is the fraction of a cluster's assigned cells. The bar above marks each subclass as one of the nineteen maternal-circuit subclasses of A–C, as detected here but outside that set, or as not detected; the two blues are nested, since the nineteen are a subset of the 84 subclasses detected. The gray strip at right gives the fraction of each cluster left unassigned. Forty-four of the 66 neuronal clusters place most of their assigned cells in subclasses we detect, accounting for 64% of their assigned neurons; the rest correspond to subclasses we do not sample, in the suprachiasmatic, arcuate, paraventricular, ventromedial and lateral hypothalamic nuclei, which lie outside the sampled coronal plane. Their 21 non-neuronal clusters, 20 of which map to subclasses detected here, are omitted from the panel.

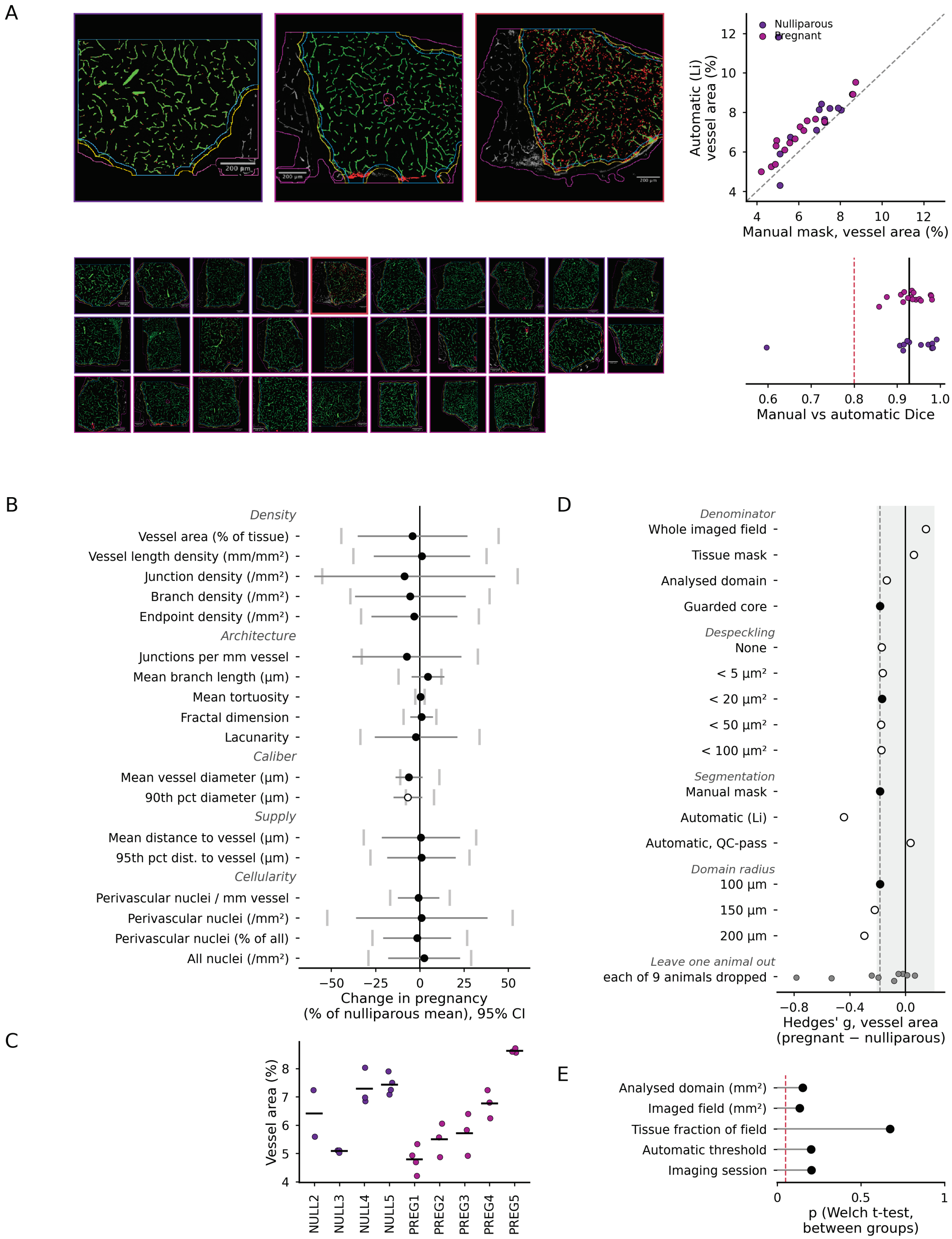

A
Nulliparous
Pregnant
Automatic (Li) vessel area (%)
Manual mask, vessel area (%)
Manual vs automatic Dice
B
Density
Vessel area (% of tissue)
Vessel length density (mm/mm²)
Junction density (/mm²)
Branch density (/mm²)
Endpoint density (/mm²)
Architecture
Junctions per mm vessel
Mean branch length (µm)
Mean tortuosity
Fractal dimension
Lacunarity
Caliber
Mean vessel diameter (µm)
90th pct diameter (µm)
Supply
Mean distance to vessel (µm)
95th pct dist. to vessel (µm)
Cellularity
Perivascular nuclei / mm vessel
Perivascular nuclei (/mm²)
Perivascular nuclei (% of all)
All nuclei (/mm²)
Change in pregnancy
(% of nulliparous mean), 95% CI
C
Vessel area (%)
NULL2
NULL3
NULL4
NULL5
PREG1
PREG2
PREG3
PREG4
PREG5
D
Denominator
Whole imaged field
Tissue mask
Analysed domain
Guarded core
Despeckling
None
< 5 µm²
< 20 µm²
< 50 µm²
< 100 µm²
Segmentation
Manual mask
Automatic (Li)
Automatic, QC-pass
Domain radius
100 µm
150 µm
200 µm
Leave one animal out
each of 9 animals dropped
Hedges' g, vessel area
(pregnant − nulliparous)
E
Analysed domain (mm²)
Imaged field (mm²)
Tissue fraction of field
Automatic threshold
Imaging session
p (Welch t-test,
between groups)

**Supplementary Fig. 7.** ***Quality control and robustness of the CD31 vascular morphometry.*** Vascular structure was quantified in an independent cohort of nulliparous and late-pregnant mice (embryonic day 17.5 to 18.5; 4 and 5 animals, 12 and 16 imaged fields of the preoptic area). Fields within an animal are technical replicates and were averaged before testing, so the unit of analysis throughout is the animal. Nulliparous, purple; pregnant, magenta. **A)** Segmentation. Top left: CD31 overlays for a typical nulliparous field, a typical pregnant field, and the field with the weakest manual-versus-automatic agreement (left to right); typical denotes the animal closest to its condition median in vessel area, and that animal's median field. In each overlay, CD31 signal is gray, the manually thresholded vessel mask green, pixels recovered only by the automatic threshold red, the imaged tissue magenta, the analyzed domain yellow and the guarded core cyan; border color gives condition, and a red border marks the one field below the Dice threshold. Bottom left: the same overlay for every analyzed field. Top right: vessel area fraction scored on the manual mask against an operator-independent Li threshold of the same fields, with the line of identity dashed (Spearman $\rho = 0.79$). Bottom right: per-field Dice coefficient between the two segmentations (black line, mean 0.93; red dashed line, the 0.80 quality-control threshold; range 0.60 to 0.99, one field below threshold). **B)** All 18 morphometric measures, expressed as the change in pregnancy as a percentage of the nulliparous mean with its 95% confidence interval, grouped into the five families labeled at left. Gray caps give the smallest change each measure had 80% power to detect. The open marker denotes the one measure quantized on the pixel grid (90th percentile diameter, 5 distinct values across 28 fields), whose p-values are tie-dominated and are not interpretable. No measure differed after correction for multiple testing (minimum FDR = 0.91). **C)** Vessel area fraction for every field, grouped by animal (points) with the animal mean (bar). The between-animal intraclass correlation is 0.85, so additional fields within an animal add little information. **D)** Effect size (Hedges' g) for vessel area fraction under each denominator, despeckling threshold, segmentation and analyzed-domain radius, and with each animal removed in turn. Filled markers are the settings used in the reported analysis; the dashed line gives the reported effect and the shaded band $|g| < 0.2$. **E)** Difference between groups in five technical covariates (Welch t-test; red dashed line, $p = 0.05$). None differed (P = 0.13 to 0.67).
